# Origin and adult renewal of the gut lacteal musculature from villus myofibroblasts

**DOI:** 10.1101/2023.01.19.523242

**Authors:** Bhargav D. Sanketi, Madhav Mantri, Liqing Huang, Mohammad A. Tavallaei, Shing Hu, Michael F. Z. Wang, Iwijn De Vlaminck, Natasza A. Kurpios

**Affiliations:** Department of Molecular Medicine, College of Veterinary Medicine, Cornell University; Ithaca, NY 14853, USA; Department of Biomedical Engineering, Cornell University; Ithaca, NY 14850, USA

**Keywords:** Single-cell RNA-seq, intestine, myofibroblast, lineage tracing, Notch signaling, smooth muscle, gut lymphatics, fat absorption

## Abstract

Intestinal smooth muscles are the workhorse of the digestive system. Inside the millions of finger-like intestinal projections called villi, strands of smooth muscle cells contract to propel absorbed dietary fats through the adjacent lymphatic vessel, called the lacteal, sending fats into the blood circulation for energy production. Despite this vital function, how villus smooth muscles form, how they assemble alongside lacteals, and how they repair throughout life remain unknown. Here we combine single-cell RNA sequencing of the mouse intestine with quantitative lineage tracing to reveal the mechanisms of formation and differentiation of villus smooth muscle cells. Within the highly regenerative villus, we uncover a local hierarchy of subepithelial fibroblast progenitors that progress to become mature smooth muscle fibers, via an intermediate contractile myofibroblast-like phenotype. This continuum persists in the adult intestine as the major source of renewal of villus smooth muscle cells during adult life. We further found that the NOTCH3-DLL4 signaling axis governs the assembly of villus smooth muscles alongside their adjacent lacteal, and we show that this is necessary for gut absorptive function. Overall, our data shed light on the genesis of a poorly defined class of intestinal smooth muscle and pave the way for new opportunities to accelerate recovery of digestive function by stimulating muscle repair.

## Introduction

The mammalian intestine is a self-renewing organ that processes ingested food, absorbing nutrients from a sea of biochemical, mechanical, and pathogenic insults. Absorption is driven by highly coordinated muscular contractions, which rely on spatiotemporal patterning and functional heterogeneity of intestinal smooth muscle (SM) cells ^1–3^. For fatty nutrients, specialized SM fibers in the intestinal villus contract to propel dietary fats through the central lymphatic capillary, known as the lacteal (**Figure 1A**) ^1,2,4–8^. Whereas most lymphatic capillaries in the body lack SM coverage, the abundance of SM fibers surrounding the lacteal is a unique feature of the contractile villus.

**Figure 1.**
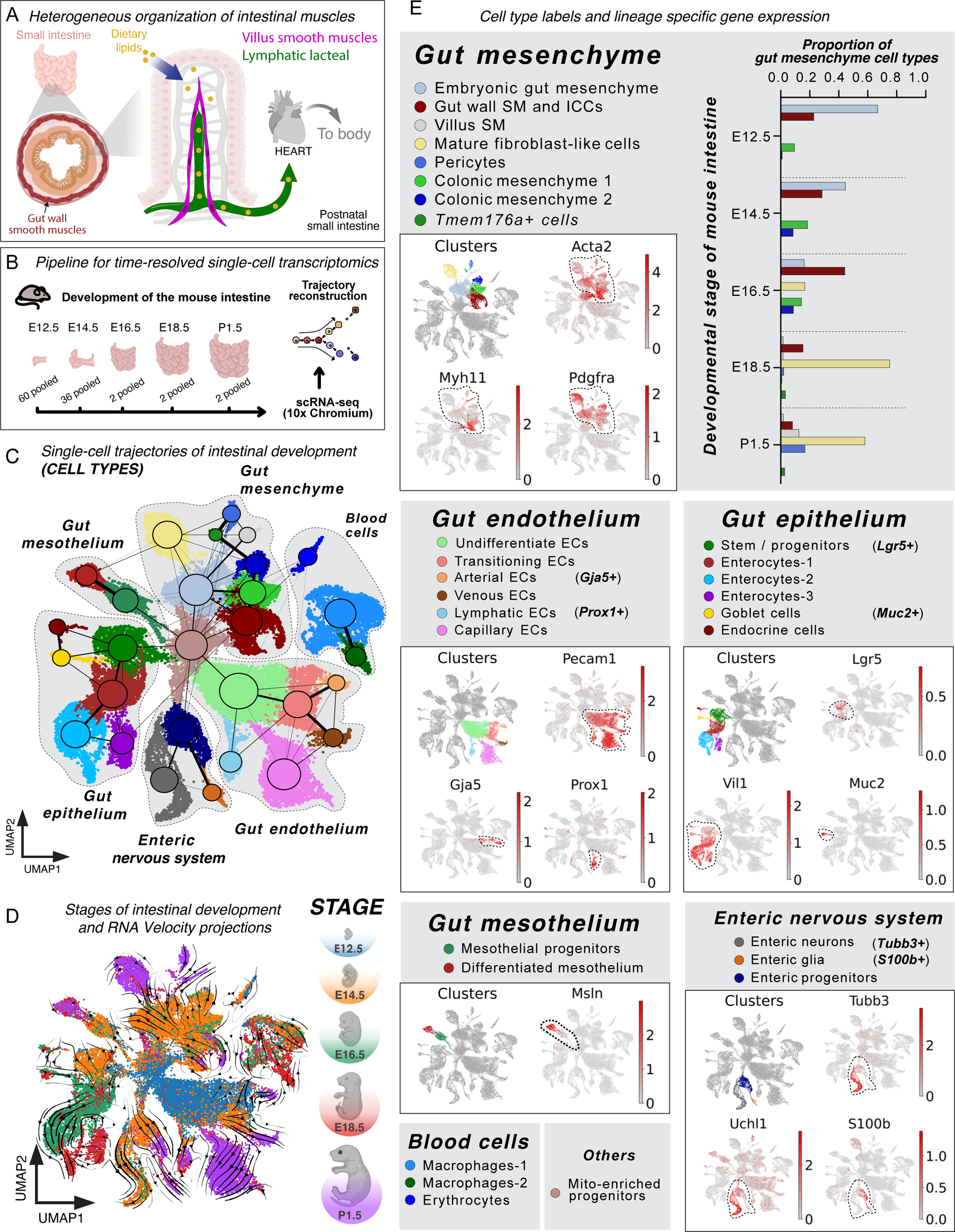
A time-resolved single-cell transcriptomic atlas of mouse intestinal development. **(A)** Cartoon depicting the distinct patterning of smooth muscles (SMs) in the gut wall and the villus. The villus contains a fat-absorptive complex of lymphatic lacteals and their associated SMs, which transport dietary lipids into the systemic circulation. (B) Experimental workflow and analysis for single-cell RNA-sequencing (scRNA-seq) at 5 embryonic and postnatal stages (E12.5, E14.5, E16.5, E18.5, P1.5). **(C)** Uniform manifold approximation and projection (UMAP) plot of 37,277 single-cell transcriptomes from mouse intestine at 5 stages, clustered by gene expression and colored by cell type labels. Partition-based graph abstraction (PAGA) map overlaid on UMAP with circles/nodes of the graph representing cell types and edges of the graph representing global relationships between cell types. The thickness of the edges in the graph is a statistical measure of connectivity between cell types, which is representative of the transcriptional similarity between them. The unsupervised mito-enriched progenitor cluster contains cells with low transcriptional diversity due to increased representation of mitochondrial genes. **(D)** UMAP plot showing mouse developmental stage of origin and transcriptomic velocity for the intestinal scRNA-seq data. **(E)** Cell type labels are segregated based on intestinal cell lineages and UMAP plots show the expression of selected clusters and cell type markers. Dotted lines on UMAP plots encircle transcriptome expression. The bar plot in the top panel shows the proportion of cells among various cell types within the gut musculature lineage at different developmental stages.

Villus SM cells were first identified in 1909 ^9^ and were subsequently shown to drive villus contractions that intensify in response to dietary fats ^10,11^. More recent *in vivo* imaging in mice has revealed that villus SM cells facilitate lipid transport by squeezing the adjacent lacteal ^2^. In addition to their contractile function, a subset of villus SM cells secretes VEGFC, the critical lymphatic morphogen. Inducible loss of either *VegfC* or its lacteal-associated receptor *Vegfr3* leads to lacteal regression and impaired dietary fat uptake ^12^.

Unlike most quiescent adult cells, gut villus lacteals and their host intestinal epithelium are in a continuous state of renewal throughout adulthood ^13,14^. This unique feature of the villus reflects the need to self-repair because the gut is constantly challenged by invasive bacteria, chemical agents, and physical constraints of peristalsis ^1,13^. Despite the crucial interplay between SM and lymphatics within the highly regenerative villus, it remains undefined how villus SM cells form, how they assemble alongside lacteals, and how they self-renew to maintain organ homeostasis throughout adult life.

To study these mechanisms, we built a single-cell gene expression atlas of the developing and postnatal mouse intestine across key developmental stages. We reconstructed the transcriptional lineage of intestinal musculature to reveal molecular heterogeneity within intestinal SM cells that are reflective of their distinct anatomical location and function, providing novel molecular markers for their isolation and discrimination. Using inducible lineage tracing, we quantified the dynamics of villus SM differentiation and uncovered a local hierarchy of subepithelial fibroblast and myofibroblast intermediates, characterized by expression of PDGFRα (platelet-derived growth factor receptor alpha) and TNC (Tenascin C), which drive the formation of SM fibers in the perinatal and adult intestine in a process that resembles wound repair ^15^. We find NOTCH3 specifically expressed along this precursor continuum with its receptor DLL4 restricted to the villus capillary vasculature. Using genetic perturbations in mice, we demonstrate that this NOTCH3-DLL4 signaling axis is a contact-dependent mechanism that governs the morphogenesis of the muscular-lacteal complex, which ultimately affects muscular contractility and lipid absorption. Collectively, our data significantly advance our understanding of the origin and renewal of the muscular-lacteal complex, a system that ensures nutrition throughout life and prevents debilitating digestive and metabolic disorders.

## Results

### A single-cell RNA-sequencing atlas of the mouse intestinal musculature

To understand the heterogeneity of diverse cell types within the developing intestinal musculature we performed single-cell RNA sequencing (scRNA-seq) to profile the mouse intestine across key developmental stages: embryonic days (E) 12.5, 14.5, 16.5, 18.5, and postnatal day (P) 1.5 (**Figure 1B**). This time window corresponds to the major transformation of the embryonic gut tube into a functional intestine ready to process fats from maternal milk at birth. We clustered 37,277 high-quality single-cell transcriptomes and used differential gene expression analysis (DGEA) and canonical markers to assign cell-type labels (**Figures 1C-1E, Figures S1A and S1B, and Table 1**). Our data reveal distinct developmental trajectories for progenitor, transitioning, and differentiated cells across all major lineages including intestinal mesenchyme, endothelium, epithelium, mesothelium, enteric nervous system, and circulating blood and immune cells (**Figures 1C and 1E, and Figure S1D**). Across the distinct lineage clusters, we calculated RNA velocity ^16^ by quantifying the proportion of molecules derived from spliced and unspliced transcripts (**Figure S1C**). RNA velocity analyses revealed trajectories from early embryonic stage E12.5 to postnatal stage P1.5 (**Figure 1D**), with evidence of progressive differentiation along each main lineage from progenitor cell states at earlier developmental stages, toward mature cells at later time points (**Figure 1E-**top-right panel, **and Figures S1D and S1E**).

To reconstruct the developmental trajectory of intestinal musculature, we specifically re-clustered 9,445 mesenchymal cells from the scRNA-seq data to further understand their transcriptional heterogeneity (**Figures S2A and S2B**). To focus our analyses on the absorptive small intestine, we excluded large intestine cells expressing *Hoxa9*, *Hoxd9*, *Colec10*, and *Adamdec1* (**Figure S2C**) ^17,18^. This approach resulted in 7,519 cells across distinct groups spanning mesenchymal progenitors to mature SM cells across all developmental stages (**Figures 2A-2D, Figure S2D and Table 2**). Within these groups, most cells from E12.5 and E14.5 were distributed among four mesenchymal progenitor clusters (MP-1, MP-2, MP-3, and MP-4) (**Figures 2A and 2B**). We further found three clusters of mature SM cells (SM-1, SM-2, and SM-3) that expressed canonical SM markers - Myosin heavy chain 11 (*Myh11*), alpha SM actin (gene: *Acta2^high^*/protein nomenclature: SMA), and Transgelin (*Tagln^high^*) (**Figures 2A and 2B**)^8,19–22^.

**Figure 2.**
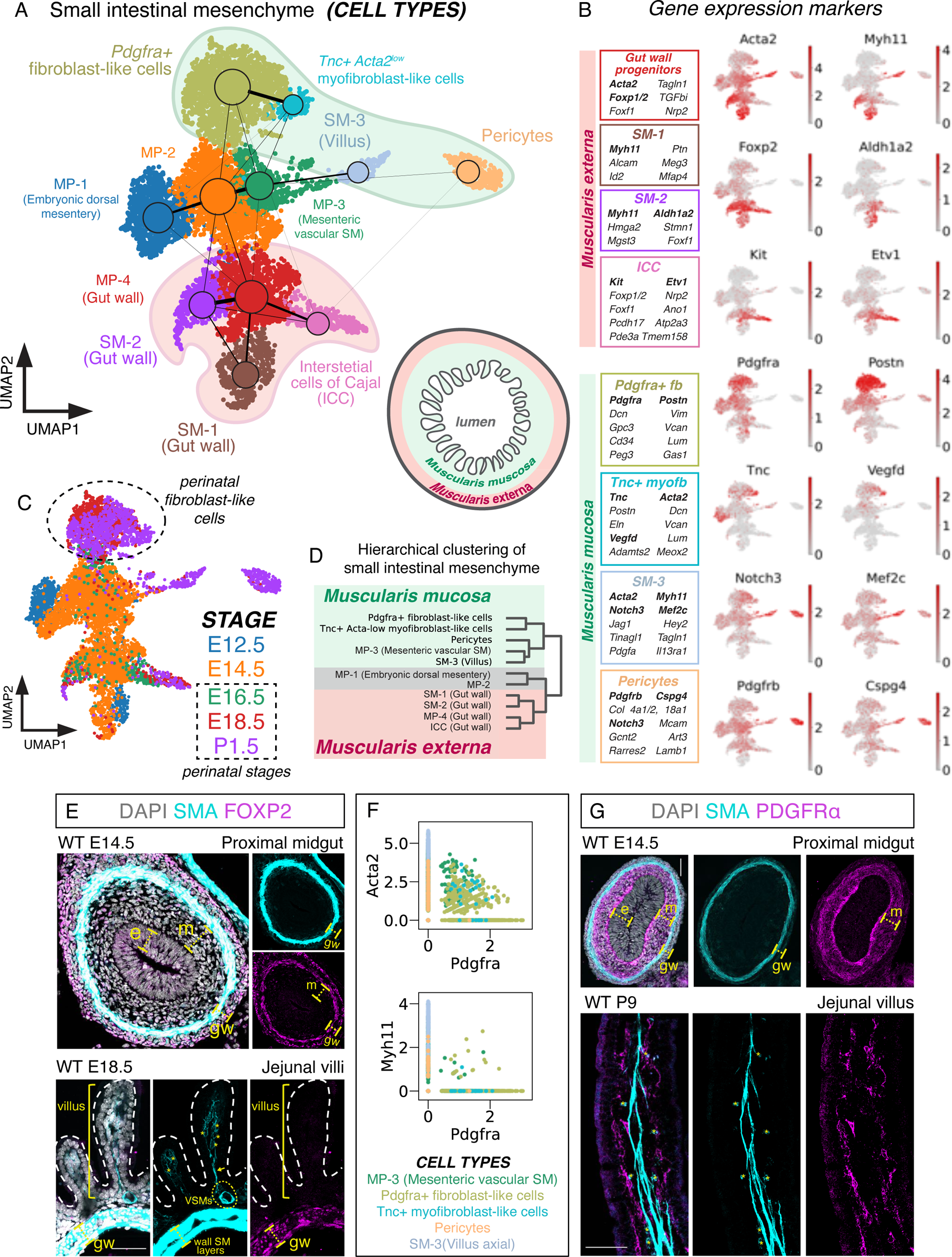
Reconstructing the developmental trajectories and transcriptional signatures of intestinal musculature. **(A)** UMAP plot of 7,519 single-cell transcriptomes from the developing small intestinal musculature at E12.5, E14.5, E16.5, and E18.5 and P1.5, clustered by gene expression and colored by cell type labels. The cartoon highlights the relative regions of assembly of the *muscularis externa* and the *muscularis mucosa* within the developing small intestine. PAGA map highlighting the transcriptional relationships between musculature cells is overlaid on the UMAP. **(B)** Boxes colored by cell type labels and UMAP plots showing the expression of selected gene markers (in bold) in the gut musculature scRNA-seq data. **(C)** UMAP plots showing mouse development stages for the small intestinal musculature scRNA-seq data. **(D)** Dendrogram showing hierarchical clustering of mesenchymal cell types in developing gut musculature. **(E)** Immunofluorescent staining of tissue sections for FOXP2 (nuclear) and SMA (cytoplasmic). FOXP2 expression colocalizes with nuclei of gut wall muscles but not SM in the proximal midgut (E14.5) and jejunum (E18.5). **(F)** Scatter plots showing co-expression of *Pdgfra* with SM markers *Acta2* and *Myh11* in cells from the *muscularis mucosa*. Cell type labels are based on Figure 2, A and B. **(G)** Immunofluorescent staining of tissue sections for PDGFRLJ (on cell membrane) and SMA (cytoplasmic). PDGFRLJ expression was observed in the mesenchymal cells between the endoderm and inner SMA+ SM layer in the proximal midgut at E14.5 and jejunal villus subepithelial mesenchyme at P9. PDGFRLJ was not expressed in SMA^high^ villus SM at P9. e - endoderm, m - gut tube mesenchyme, gw - gut wall, asterisk (*) - star cell, arrow - spindle-shaped villus SM cell, VSMs - vascular SMs. All scale bars = 50μm.

Mature SM cells of the small intestine include a heterogeneous mix of villus SM within the *muscularis mucosa* and the two layers of circular and longitudinal SM of the *muscularis externa* (gut wall) (**Figure 2A**) ^23^. These populations differentiate radially in sequence during intestinal development, with the outer circular SM layer developing first, followed by the longitudinal SM layer, and finally, the villus SM fibers arising with lacteal formation close to birth ^8,24^. This sequence was evident within our unsupervised clustering, with each of the three SM clusters exhibiting unique transcriptional signatures (**Figures 2A, 2B, and 2D, and Figure S2E**), and distinct developmental stage composition (**Figures 2A and 2C**). SM-1 and SM-2 contained cells from E12.5 and E14.5 (**Figures 2A and 2C**), consistent with the stepwise differentiation pattern of the emerging gut wall ^8^. In contrast, SM-3 contained cells mostly from P1.5, indicating villus SM identity (**Figures 2A and 2C**) ^24^. Several findings further support these classifications. In a cluster along the trajectory of gut wall progenitors of SM-1 and SM-2, but not SM-3, we found an enrichment of *c-Kit* and other markers of Interstitial Cells of Cajal (ICC) ^1,25^, which are known to be spatially restricted to the *muscularis externa* (**Figures 2A and 2B**). We also identified Forkhead box protein p2 (*Foxp2*) and Neuropilin 2 (*Nrp2*) transcripts to be differentially expressed in the *muscularis externa* clusters (**Figure 2B and Figure S3A**).

Validating the spatial fidelity of our cell type classifications in mouse intestinal tissue, we found nuclear-localized FOXP2 protein expression in SMA+ cells of the gut wall but not in the villus (**Figure 2E**), consistent with the notion that SM-1 and SM-2 represent gut wall musculature and MP-4 represents their mesenchymal progenitors. Immunofluorescence also confirmed NRP2 expression in the *muscularis externa*, in addition to its previously described expression in lymphatic endothelial cells (**Figure S2F**) ^26^. To further differentiate the molecular regulation of cellular diversity within the developing mesenchyme, we performed transcriptional network analysis using SCENIC (**Figures S3A and S3B**) ^27^. Together, our transcriptomic analysis captured the differentiation of gut mesenchyme into the diverse musculature of the gut wall and the villus stroma.

### Villus SM cells arise from SMA^low^ star cells by PDGFR**α**+ fibroblast-to-myofibroblast transition

Platelet-derived growth factor receptor alpha (PDGFRα) has recently emerged as a marker of mesenchymal stem and progenitor cells across multiple organ systems ^28^. During villus formation, PDGFRα+ mesenchymal cells accumulate beneath the endoderm forming a subepithelial fibroblast cell population (**Figure S4A**) ^29–35^. In our transcriptomes from the intestinal musculature, we found *Pdgfra* expression in mesenchymal progenitor clusters at E12.5 and 14.5, and in a fibroblast-like cluster from later (perinatal) time points (E16.5, E18.5, and P1.5) marked by Periostin (*Postn^high^*) (**Figures 2A-2C**). In contrast, *Pdgfra* was absent in mature SMs, ICCs, and a pericyte cell cluster marked by *Cspg4* and *Pdgfrb^high^*(**Figures 2A and 2B**) ^22^. Using two-dimensional (2D) scatter plots of our transcriptomes from the *muscularis mucosa*, we observed a stark inverse relationship between the expression of *Pdgfra* and *Acta2*, while cells expressing *Pdgfra* and *Myh11* were almost mutually exclusive (**Figure 2F**). Interestingly, whereas the *Pdgfra+* fibroblast-like cluster was broadly negative for *Acta2*, we noted a subset of cells that were *Acta2^low^* (**Figures 2A and 2B**), suggesting a cell state transition from fibroblast to myofibroblast ^19,36–38^.

During physiological wound healing and pathological fibrosis, fibroblast cells progress through a precursor stage to become contractile myofibroblasts, by gradually expressing SMA, arranged as stress fibers ^36,39,40^. In our previous studies of the villus SM ^24^, we detected a subset of SMA^low^ villus mesenchymal cells with stellate morphology, which we called star cells, and hypothesized that they represent a precursor state for villus SM differentiation in the embryonic and postnatal intestine. Using immunofluorescence to validate these expression patterns, we found PDGFRα protein in undifferentiated mesenchymal cells just beneath the gut endoderm at E12.5 and E14.5 (**Figure 2G and Figure S4C**). At E18.5 and P9, immunofluorescence confirmed PDGFRα expression in the villus subepithelial cells that were closely positioned to, but distinct from the mature (SMA^high^) SM cells, whereas SMA^low^ star cells displayed variable levels of PDGFRα expression (**Figure 2G and Figure S4C**). We also analyzed mice with dual genetic reporters - H2b-GFP (nuclear green fluorescent protein) under the control of the endogenous *Pdgfra* gene promoter ^41^ and dsRed (red fluorescent protein) under the control of a transgenic *Acta2* promoter ^42^. Here, whereas we observed H2b-GFP+ cells in both the villus stroma (GFP high and low) and the sub-mucosal regions (GFP low), dsRed+ cells were mostly limited to the villus stroma (**Figure 3A and Video S1**), suggesting an increased presence of myofibroblast-like transition within the villus during SM differentiation. Whereas star-shaped cells showed variable levels of H2b-GFP and dsRed expression, spindle-shaped SM cells were mostly dsRed+ and H2bGFP- (**Figure 3A and Video S1**), consistent with our PDGFRA and SMA protein expression (**Figure 2G and Figure S4C**).

**Figure 3.**
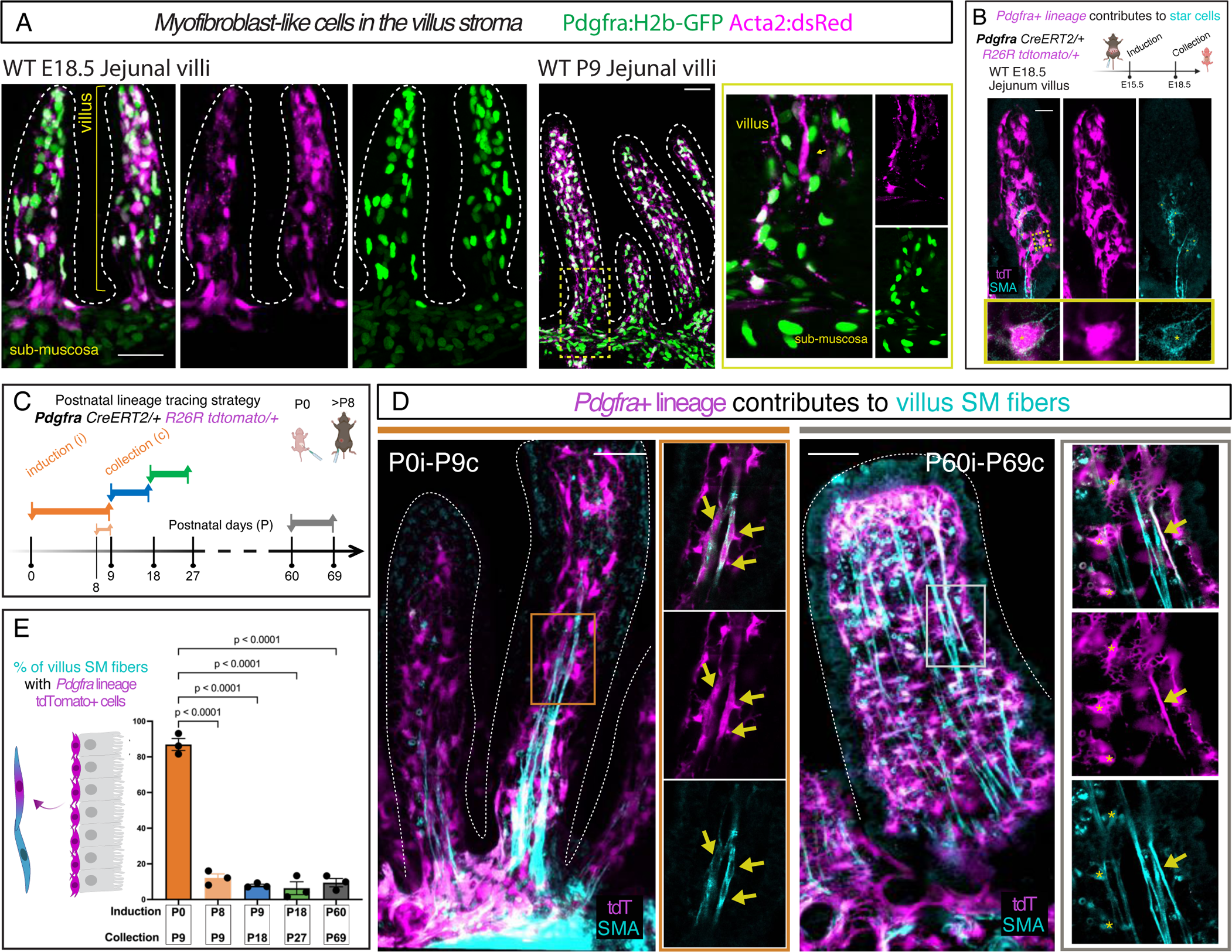
Pdgfra+ villus fibroblast-like cells differentiate to assemble villus SM. **(A)** E18.5 and P9 jejunal villi visualized with genetic reporters *Pdgfra: H2b-GFP* and *Acta2:dsRed*. **(B)** Induction of lineage tracing in *Pdgfra+* cells of pregnant mouse dams at E16.5 results in tdTomato reporter (intracellular) labeling of SMA^low^ star cells (yellow asterisks) in the villus of the mouse pup at E18.5. The yellow box shows a magnified inset of a tdTomato+ SMA^low^ star cell. **(C)** Schematic for *Pdgfra+* lineage tracing in 5 induction-collection intervals of P0i-P9c, P8i-P9c P9i-P18c, P18i-P27c, and P60i-69c. **(D)** *Pdgfra+* lineage tracing in induction-collection intervals P0i-P9c and P60i-69c; whole-mount immunostaining of SMA on jejunal villi. Boxes (orange and gray) show the region magnified in the inset with a single 1μm section along the whole-mount z stack. Yellow arrows point to SMA+ fibers with tdTomato expression**. (E)** Quantification of *Pdgfra+* lineage tracing contributions to SM fibers in induction-collection intervals P0i-P9c, P8i-P9c P9i-P18c, P18i-P27c, and P60i-69c. Comparisons by one-way ANOVA followed by Tukey’s multiple comparisons test (presented as mean ± SEM). tdT - tdTomato reporter. All scale bars = 50μm.

Based on these patterns, we hypothesized that during late embryogenesis, *Pdgfra+* fibroblast-to-myofibroblast transition represents a hierarchical continuum of stem and progenitor cell states involved in villus SM differentiation. To test this, we performed *Pdgfra+* lineage tracing using the *Pdgfra-CreERT2* driver ^43^ and *Rosa26-tdTomato* reporter mice ^44^ prior to the formation of star cells (E15.5) (**Figure 3B**). This resulted in *tdTomato* labeling of SMA^low^ star cells in the villus stroma at E18.5 (**Figure 3B**). Since villus SM are mostly assembled postnatally, we performed *Pdgfra+* lineage tracing at multiple postnatal induction (i) - collection (c) intervals and quantified the contribution of the *Pdgfra+* lineage to the formation of mature SM fibers (**Figure 3C**). We observed *Pdgfra+* lineage reporter *tdTomato* expression in 87±3.4 % of villus SM fibers in the P0i-P9c interval (**Figures 3D and 3E**, **and Video S2**). In P8i-P9c (1-day interval), although we observed similar and extensive labeling of the villus fibroblasts, SM fibers were sparsely labeled (12±2.3 %) with *tdTomato*+ cells (**Figure 3E and Figure S5A**). This indicates that the extensive SM contribution observed in the P0i-P9c interval resulted from the differentiation of *tdTomato*+ fibroblasts. In the next induction intervals tested (neonatal - P9i-P18c and P18i-P27c, and adult - P60i-69c, **Figure 3C**), the *Pdgfra+* lineage sparsely labeled 7.9±0.72 %, 6.3±3.6 % and 9.4±2.5% of the SM fibers, respectively (**Figures 3D and 3E**, **Figure S5A, and Video S3**). Altogether, these data suggest that during mouse perinatal development and adult homeostasis, *Pdgfra+* fibroblasts progress to become SMA^low^ myofibroblast-like cells that differentiate into villus SM cells.

### Villus SM cells renew from TNC+ intestinal myofibroblasts

The transition from fibroblasts to contractile myofibroblasts is paramount to organogenesis, tissue repair, and cancer ^15^ but the mechanisms governing this transition *in vivo* are poorly understood. To investigate the transcriptional changes during the formation of SMA^low^ star cells (a fibroblast-to-myofibroblast transition), we performed unsupervised clustering of the 1,708 perinatal fibroblast-like cells we had captured (**Figure 2C**). This partitioned them into four groups based on their transcriptomic programs (**Figure S6A**). Interestingly, the gene expression signatures previously associated with mature fibroblast diversity in the adult villus ^22,35,45^ could not account for the results of our unsupervised clustering of perinatal fibroblasts (**Figures S6B-S6D**). Instead, we observed the expression of previously reported markers of myofibroblasts within these clusters ^19,21,21,36,39,41,42^, including Tenascin-c (*Tnc*), which exhibited expression patterns most akin to *Acta2* and *Tagln* (**Figure S6E**). The *Acta2+ Tagln+ Tnc+* cells exhibited expression of Hedgehog signaling effectors *Ptch1/2*, *Gli1*, and *Hhip* (**Figure S6F**), consistent with prior findings that alterations in Hedgehog signaling within the intestinal epithelium could either enhance or diminish SM differentiation ^46^.

TNC is an extracellular matrix protein known to induce actin stress fibers during fibroblast-to-myofibroblast transition ^47,48^ and its expression has been previously detected in the subepithelial fibroblasts of the adult villus ^14,49,50^. In our scRNA-seq analysis, 2D scatter plots revealed that *Tnc* expression is low in *Pdgfra^high^*perinatal fibroblasts but peaks in *Pdgfra^low^* and *Acta2^low^*myofibroblast intermediates, commensurate with the appearance of the SMA^low^ star cells in the villus ^24^, while mature SM cells expressing Myh11 are rarely Tnc+ (**Figure 4A**). Immunofluorescence validated TNC protein expression in SMA^low^ cells at E18.5 and P9, and its absence in mature SM fibers (**Figure 4B and Figure S7A**). In our scRNA-2seq analysis, *Tnc* was also expressed in the MP-1 cluster composed of mesenchymal cells from E12.5 and E14.5 (**Figure 2A and Figure S7B**). We confirmed these results and observed that TNC expression during these early stages is localized to the mesenchymal cells of the embryonic dorsal mesentery, identified by the embryonic left-right asymmetry marker *Pitx2* ^51,52^ (**Figure S7B**). Importantly, these TNC+ cells are distinct from the villus myofibroblasts that first appear perinatally ^24^. Together, these data suggest that during villus SM differentiation, *Pdgfra+* fibroblast progenitors transition through a *Tnc+*/*Acta2^low^* myofibroblast precursor cell state, equivalent to star cells we have previously described *in vivo* ^24^.

**Figure 4.**
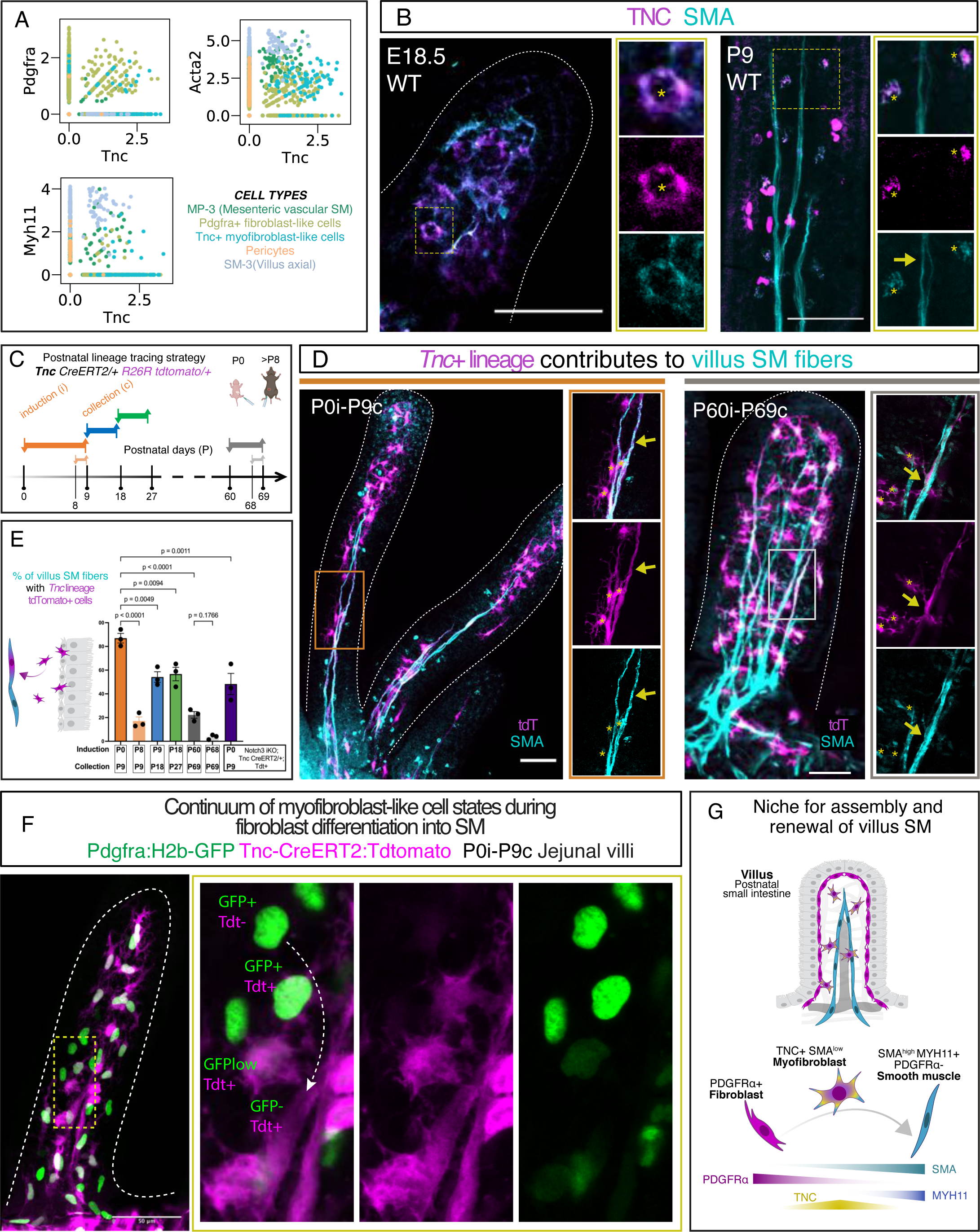
Villus SM arises from fibroblasts that transition through a continuum of Tnc+ myofibroblast-like cell states. **(A)** Scatter plots showing co-expression of *Tnc* with *Pdgfra, Acta2*, and *Myh11* in the cell types of the developing *muscularis mucosa*. **(B)** Immunofluorescent staining of TNC and SMA on jejunal villi at E18.5 and P9. TNC expression (intra/extracellular) colocalizes with the SMA^low^ star cells but not with the SMA^high^ villus SM. Yellow boxes show the region magnified in the inset. In the P9 insets, yellow asterisks mark SMA+ TNC+ star cells whereas the yellow arrow marks a spindle-shaped SM. **(C)** Schematic for *Tnc*+ lineage tracing in 6 induction-collection intervals of P0i-P9c, P8i-P9c P9i-P18c, P18i-P27c, P60i-69c, and P68i-P69c. **(D)** *Tnc*+ lineage tracing in induction-collection intervals P0i-P9c and P60i-69c; whole-mount immunostaining of SMA of jejunal villi. Boxes (orange and gray) show the region magnified in the inset with a single 1μm section along the whole-mount z stack. Yellow arrows point to SMA+ fibers with tdTomato expression. Yellow asterisks mark *Tnc*+ lineage myofibroblasts. **(E)** Quantification of *Tnc*+ lineage tracing contributions to SM fibers in induction-collection intervals P0i-P9c, P8i-P9c P9i-P18c, P18i-P27c, P60i-69c, P68i-P69c, and for Notch3 iKO at P0i-P9c. Comparisons by one-way ANOVA followed by Tukey’s multiple comparisons test (presented as mean ± SEM). **(F)** Jejunal villi visualized with genetic reporter *Pdgfra: H2b-GFP* and *Tnc*+ lineage tracing at P0i-P9c interval. A continuum of states can be visualized based on GFP and tdTomato expression. The white arrow denotes the proposed trajectory of fibroblast to SM transition through a myofibroblast intermediate. (G) Proposed model of fibroblast to SM transition through a myofibroblast intermediate in the intestinal villus. Intermediate cells fall along the continuum of gene expression of SMA (*Acta2*), MYH11, PDGFRα, and TNC. tdT - tdTomato reporter. All scale bars = 50μm.

To assess the fate of *Tnc+* cells in the postnatal stages, we performed lineage tracing using the *Tnc-CreERT2* driver ^53^ at fixed induction-collection intervals, as above (**Figure 4C**). *TdTomato* expression was observed in villus myofibroblast-like cells (asterisks) and in 87±4%, 54±4.5%, 57±5.7% and 22.3±2% of SM fibers (arrows) in the intervals P0i-P9c (starting at birth), P9i-P18c and P18i-P27c (neonatal), and P60i-69c (adult), respectively (**Figures 4D and 4E**, **Figure S7C, Video S4, and Video S5**). At 1-day intervals (P8i-P9c and P68i-P69c), whereas *tdTomato*+ myofibroblasts were similarly abundant, *tdTomato+* SM fibers were less common during these short intervals and were mostly identified as transitioning SM cells integrating into axial fibers (17±3.5% for P0i-P9c; 3.6±1.2% for P68i-P69c) (**Figure 4E**, **Figures S7D and S7E, Video S6 and Video S7**). These data confirm that the *tdTomato*+ SM fibers observed in longer intervals such as P0i-P9c and P60i-69c arise from transitioning myofibroblasts from *Tnc+* lineage. Interestingly, whereas the contribution of *Pdgfra+* and *Tnc+* lineages to villus SM fibers was comparable at birth (P0i-P9c, 87±4% vs. 87±3.4%), the *Tnc+* lineage contribution at later induction times was more pronounced than that of the *Pdgfra+* lineage. This implies that *Tnc+* myofibroblasts continue to exist in the mature intestine, serving as a renewable pool of adult SM stem/progenitor cells.

We further examined the progression of myofibroblast cell states as fibroblasts transitioned into mature SM by tracking *Tnc+* lineage in combination with *Pdgfra* promoter-driven H2b-GFP (**Figure 4F, Video S8 and Video S9**). This tracking approach enabled us to visualize the complete continuum of transitions from fibroblast to myofibroblast to mature SM fibers (fibroblast: H2B-GFP+*tdTomato-*, myofibroblast: H2B-GFP+*tdTomato+* or H2B-GFPlow/- *tdTomato+* and recognized by their stellate morphology, and mature SM fiber: H2B-GFPlow/- *tdTomato+* and recognized by their spindle morphology). In summary, our observations suggest that villus SM cells originate from *Tnc+* myofibroblast precursors that originate from the *Pdgfra*+ lineage (Figure 4G).

To learn whether the above features of myofibroblast-to-villus SM trajectory are conserved in the human, we examined the mesenchymal trajectories from the human gut cell atlas ^54^ (**Figure S8A**). Similar to the mouse, we found three human *MYH11+ACTA2highTAGLNhigh* SM clusters. Among them, the cluster annotated as ‘Contractile pericytes’ resembled the mouse villus SM cluster based on gene expression (NOTCH3+, MEF2C+, etc.) (**Figure S8B**). Associated clusters ‘Stromal 3’ (POSTN+PDGFRA+) and ‘Angiogenic pericytes’ (TNC+ACTA2+PDGFRA-) resembled mouse villus fibroblast and myofibroblast clusters, respectively. Taken together, these observations suggest that the molecular characteristics of the myofibroblast-to-villus SM trajectory in the mouse are also preserved in the human villus.

**NOTCH3-DLL4 signaling is necessary for muscular-lacteal assembly and fat absorption** Villus SM cells arise in parallel with the formation of lacteals (E18.5) to generate the muscular-lacteal complex ^24^. Despite its essential function in fat absorption, the mechanisms that govern the assembly of these adjacent cells remain unknown. As part of our effort to understand the molecular nature of the muscular-lacteal interaction, we noted that the juxtacrine (contact-dependent) signaling receptor *Notch3* was strongly enriched in Tnc+ myofibroblasts and villus SM cells (**Figure 4A**). In further support of active NOTCH signaling within the *muscularis mucosa*, we found multiple components of this pathway including *Jagged1*, *Hey2, Notch1* (and their counterparts) expressed in both the mouse and human datasets (**Figure S8C**) ^55^. In our 2D scatter plots, *Notch3* levels were correlated with *Acta2* and *Myh11* expression, consistent with NOTCH signaling activity along the myofibroblast-to-villus SM trajectory (**Figure 5A**). While most studies focus on the actions of NOTCH signaling in the endothelium, *Notch3* is the predominant NOTCH receptor responsible for vascular SM maturation and is the causal gene for the hereditary vascular dementia CADASIL, characterized by degeneration of SM cells ^56–58^. Through immunofluorescence at E18.5 and P9, we confirmed NOTCH3 protein expression in SMA^low^ star cells and in SMA^high^ mature SM fibers (**Figure 5B**). Furthermore, we confirmed the protein expression of Hairy/enhancer-of-split related with YRPW motif protein 2 (HEY2, **Figure S8C and Figure S9A**) and Melanoma cell adhesion molecule (MCAM, **Figure S8C, and Figure S9B**), which are known to be associated with a vascular SM phenotype ^59–61^. Cluster MP-3, mainly comprising cells from E14.5, exhibited gene expression patterns resembling villus SM and its progenitors, including genes like *Acta2*, *Myh11*, *Pdgfra*, *Tnc*, and *Notch3* (**Figure 2F**, **Figure 4A, and Figure 5A**). Of interest, a gene with differential expression within this cluster, WNT4A, was found to be specifically localized in vascular SM in the vicinity of the mesenteric blood vasculature (**Figure S9C**). These observations indicate that villus SM cells exhibit a molecular similarity to blood vessel-associated SM cells and are set apart from the SM found in the gut wall.

**Figure 5.**
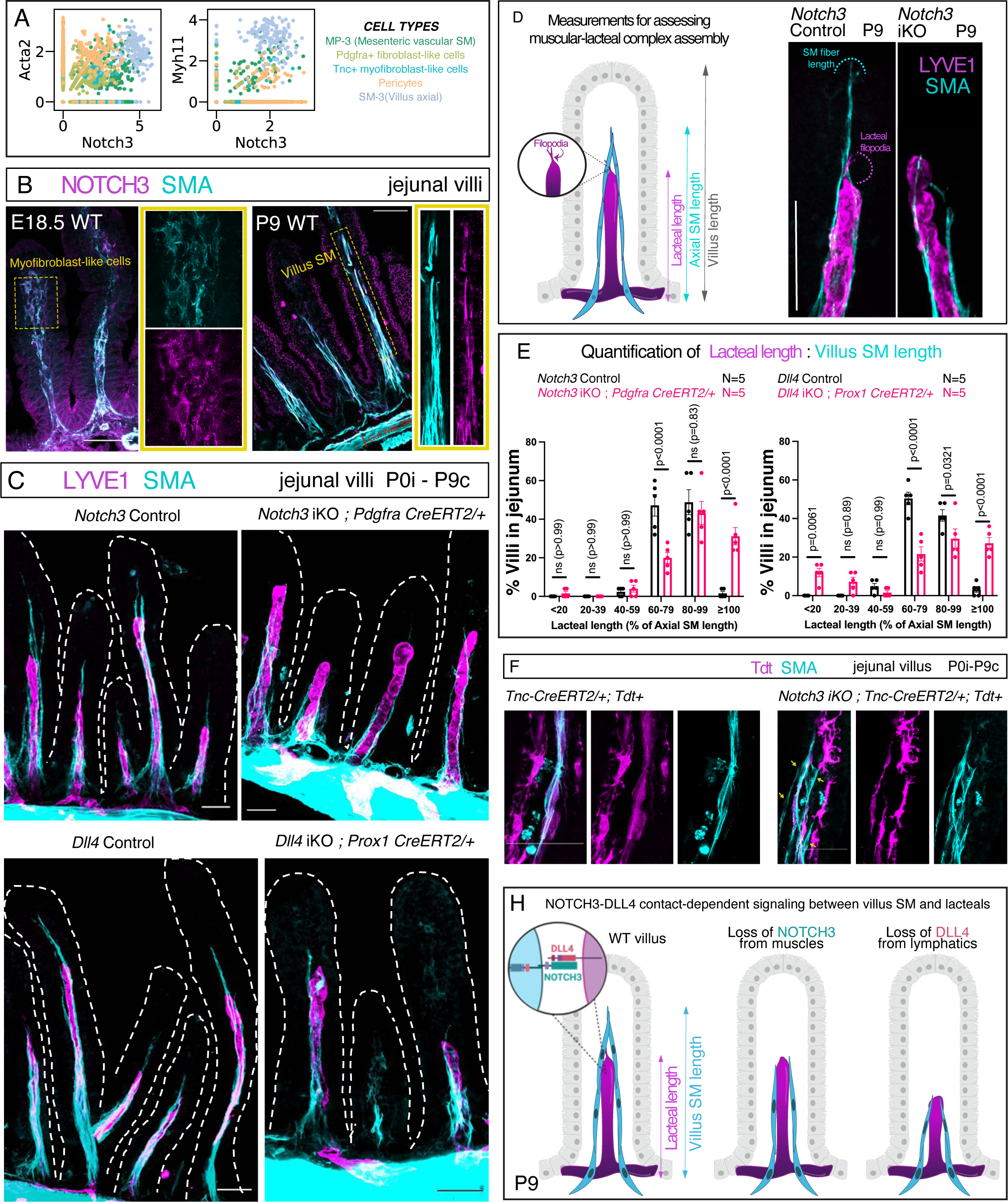
Loss of NOTCH3-DLL4 signaling perturbs villus SM assembly alongside lacteal. **(A)** Scatter plots showing co-expression of *Notch3* with SM markers Acta2 and Myh11 in the cell types of the developing *muscularis mucosa*. **(B)** Immunofluorescent staining of jejunal villi tissue sections at E18.5 and P9 for NOTCH3 and SMA expression. NOTCH3 expression (on the cell membrane) colocalizes with the SMA^low^ star cells and SMA^high^ villus SM (SMA - cytoplasmic). Yellow boxes show the region magnified in the inset. **(C)** Whole mount immunofluorescence of SMA and lymphatic marker LYVE1 on villi from the proximal jejunum upon inducing knockout of *Notch3* gene using the *Pdgfra CreERT2* driver (top panel) and *Dll4* gene knockout using *Prox1 CreERT2* (bottom panel). **(D)** Cartoon schematic for measurement of villus, lacteal, and SM lengths and filopodia presence (left). Representative loss of *Notch3* phenotype showing reduced villus SM length and lacteal filopodia (right). **(E)** Binned quantification of the percentage of villi with given ratios of lacteal and villus SM lengths upon *Notch3* deletion using *Pdgfra-CreERT2* and *Dll4* iKO using *Prox1-CreERT2*, compared using two-way ANOVA followed by Šídák’s multiple comparisons test (presented as mean ± SEM). **(F)** *<u>Tnc</u>*+ lineage tracing at P0i-P9c interval with or without Notch3. Whole-mount immunostaining of SMA in jejunal villi. Yellow arrows point to tdTomato-SM cells. **(G)** Proposed model of NOTCH3-DLL4 function during the assembly of the muscular-lacteal complex, as seen at P9. iKO - inducibly knocked out. tdT - tdTomato reporter. All scale bars = 50μm.

NOTCH is a short-range signaling pathway that occurs between membrane-bound receptors and ligands expressed on adjacent cells. Of interest, the Notch ligand Delta-like 4 (DLL4) is highly expressed in tip cells of sprouting lacteals where it directs filopodia-mediated lacteal survival and regeneration ^14^. We validated DLL4 expression in the perinatal lacteal tip cells using the lymphatic endothelial cell marker LYVE1 ^62^ and Prox1-GFP lymphatic reporter mice ^63^ (**Figures S9D and S9E**). Based on these gene expression patterns, we hypothesized that contact-dependent NOTCH3-DLL4 signaling is necessary for the assembly of the muscular-lacteal complex. To test this, we deleted *Notch3* or *Dll4* from villus SM or lymphatic endothelial cells, respectively ^64,65^. Loss of NOTCH3 and DLL4 protein upon induction of Cre was confirmed by immunostaining (**Figures S10A and S10B**). Deleting *Notch3* at P0 (with *Pdgfra-CreERT2*, inducibly knocked out - iKO) (**Figures 5C and 5D**, **and Figure S11A**) or in the embryonic gut mesenchyme (with *Hoxb6-Cre* ^66^) (**Figure S10C and Figure S11B**) resulted in a significant reduction in the length of villus SM fibers at P9. A similar reduction in the length of SM fibers was observed when Dll4 was removed from lacteals (P0, with *Prox1-CreERT2*) ^67^ (**Figure 5C and Figure S11C**). We found that SM fibers were not extended beyond their adjacent lacteal tip in 31.3% of the villi upon *Notch3* deletion using *Pdgfra-CreERT2* at P0 (**Figures 5D and 5E**, **and Video S10**), in 37.3% using *Hoxb6-cre* (**Figure S10C**), and in 27.2% when *Dll4* was deleted from lacteals (**Figure 5E**). This was statistically significant when compared to control mice where SM fibers that did not extend beyond lacteals were rarely found (<4% of villi) (**Figure 5E, Figure S10C and Video S11**). *Notch3* deletion from the musculature did not affect lacteal length (**Figures S7D and S7E**), but *Dll4* deletion from lacteals reduced lacteal length (≤ 20% villus length) with 16% of the mutant villi developing without lacteals (**Figure S7F**). However, deletion of either *Notch3* or *Dll4* resulted in a 40-50% reduction in the number of lacteal filopodia (**Figures S7G-S7I**). Inducing *Notch3* deletion with *Pdgfra-CreERT2* in the 1-day interval (P8i-P9c, which results in recombination in fibroblasts but not SM, **Figure 3E and Figure S5B**) did not significantly alter SM assembly (**Figure S12A**). This suggests that the muscular-lacteal phenotypes when *Notch3* is deleted within the P0i-P9c interval are mainly a direct effect of *Notch3* on SM, rather than indirect effects on the stroma. When *Notch3* was deleted embryonically using *Hoxb6-Cre*, but not postnatally using *Pdgfra-CreERT2*, we also observed a loss of vascular SM coverage of the mesenteric blood vasculature (**Figure S12B**). These data are consistent with the molecular resemblance between villus SM and the mesenteric vascular SM progenitors (MP-3) (**Figure 2F**, **Figure 4A**, **Figure 5A, and Figure S9C**) and underscore the significance of temporal regulation when investigating the origin and assembly of intestinal SM.

We also analyzed the effects of Notch3 deletion from myofibroblasts using *Tnc-CreERT2* and *tdTomato* reporter and observed a significant reduction in *tdTomato*+ SM fibers (48±8.9% vs. 87±4% control, **Figure 5F and Figure 4E**). These data indicate that Notch3 loss from myofibroblasts impairs their ability to develop into SM, aligning with the well-established role of NOTCH signaling in vascular SM maturation ^56–58^. However, the muscular-lacteal complex was largely intact in these mice suggesting compensatory mechanisms from fibroblasts that escaped *Notch3* deletion and/or other sources. Collectively, these results indicate that NOTCH3-DLL4 signaling is necessary for the assembly of villus SM fibers alongside their adjacent lacteals (**Figure 4H**).

To evaluate fat absorption within the muscular-lacteal complex after *Notch3* loss, we administered the fluorescent long-chain lipid tracer BODIPY C-16 ^68^ to control and *Notch3* iKO (utilizing *Pdgfra-CreERT2*) mouse pups at P9 (**Figure 6A**). Following *Notch3* deletion, we observed that BODIPY C-16 accumulated in the distal jejunal villi, while it was completely cleared from the villi of control pups (**Figure 6B**). To quantify the lipid flux, we gavaged control and *Notch3* iKO (utilizing *Pdgfra-CreERT2*) mouse pups at P9 with olive oil and measured triglycerides and cholesterol in blood plasma. While total cholesterol, high-density lipoprotein cholesterol, and low-density lipoprotein cholesterol showed no notable differences between *Notch3* iKO and control pups, there was a significant reduction in blood triglyceride concentrations in *Notch3* iKO pups compared to controls (mean 150 mg/dL vs. 251.25 mg/dL). These results indicate a reduction in triglycerides absorbed through the villus muscular-lacteal system as chylomicrons ^2,5,69^ (**Figure 6C**). In summary, these findings collectively suggest that the structural defects in the muscular-lacteal complex resulting from *Notch3* deletion functionally impair villus lipid absorption.

**Figure 6.**
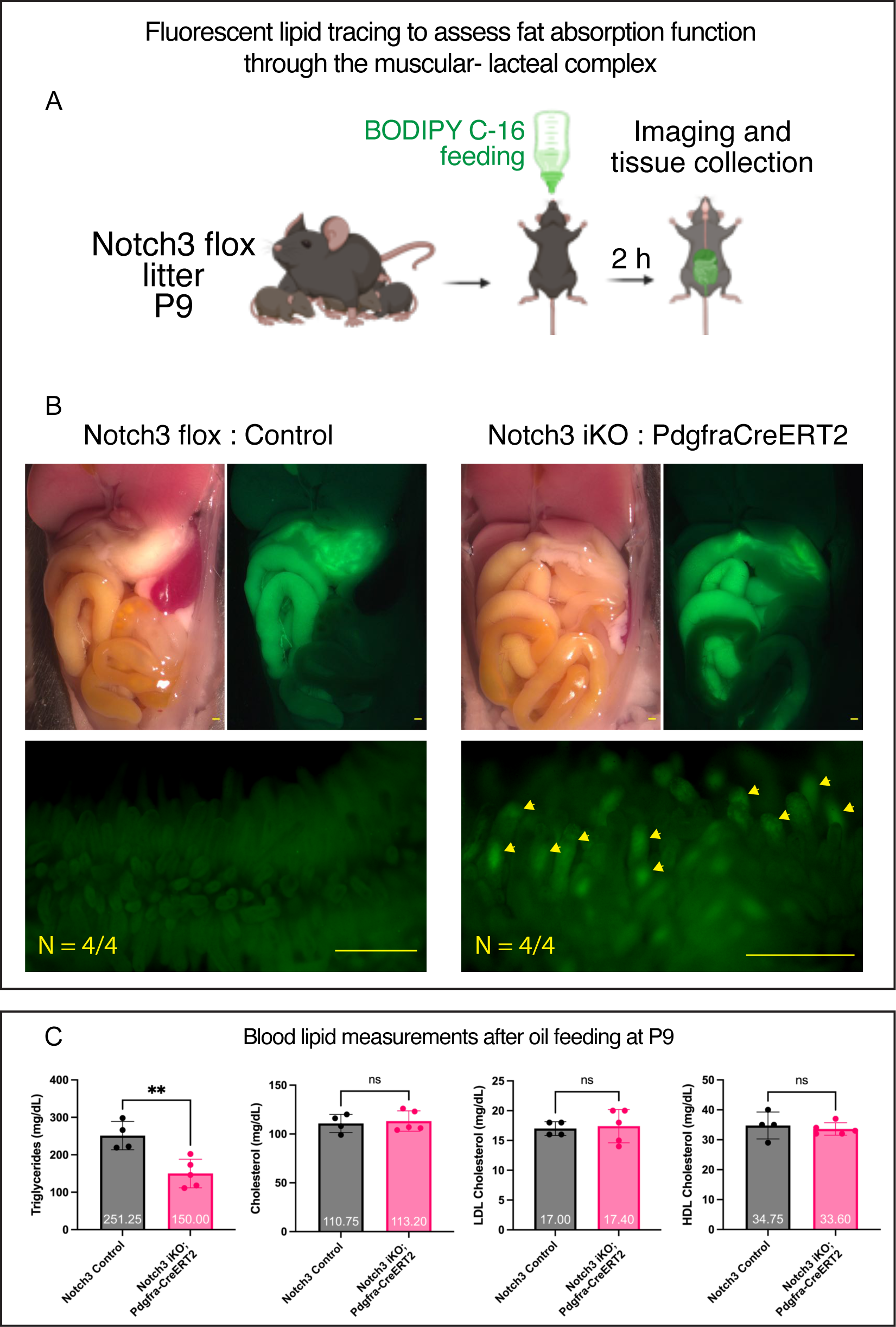
Loss of *Notch3* impairs the clearance of lipid tracer through the muscular-lacteal complex. **(A)** Schematic for feeding and imaging fluorescent lipid tracer BODIPY C-16 in mouse pups. **(B)** Fluorescent imaging of lipid tracer BODIPY C-16 in the abdominal cavity (top) and unfixed distal jejunal villi (bottom) 2 hours post-gavage. BODIPY C-16 accumulated in the villi of *Notch3* iKO but was cleared from the villi of control pups. **(C)** Measurements of triglycerides and cholesterol from control and *Notch3* iKO pups induced in the P0i-P9c interval after oil feeding. The comparison was made using an unpaired t-test (presented as mean ± SEM). All scale bars = 500μm.

## Discussion

Intestinal villi are the central units for the absorption and transport of dietary nutrients. They undergo constant regeneration to withstand the harsh conditions of the digestive tract. The coordinated movement and contractility of villi, vital for lipid transport through lymphatic lacteals, are facilitated by the patterned assembly of SM fibers alongside lacteals ^2,4,5^. Unlike other cell types in the intestine, such as the epithelium, our understanding of the stem and progenitor cell states in the gut mesenchyme and the intercellular processes that regulate the formation and renewal of villus SM has been limited by the absence of detailed gene expression data and genetic tools. To bridge this gap in knowledge, we embarked on a project to generate a time-resolved transcriptomic map of mouse intestinal development. To this end, we isolated and analyzed cells from whole organ preparations, enabling us to capture the full spectrum of cellular diversity spanning multiple intestinal cell lineages. This comprehensive approach allowed us to reconstruct the differentiation trajectory of mesenchymal cells and pinpoint the specific genetic programs responsible for generating the diverse musculature present in the gut.

Contractile activity leads to regular mechanical breakdown of muscle cells, requiring them to be renewed. For example, skeletal muscle satellite cells are adult stem cells that maintain skeletal muscle homeostasis ^70^. However, the molecular mechanisms behind the assembly and maintenance of villus SM within the intestine are not yet understood. Ultrastructural and genetic studies have detected distinctly shaped mesenchymal cell variants at different proximities from villus SM fibers ^24,71–73^. Recent research has revealed the crucial signaling role of villus mesenchymal cells in the zonation, maintenance, and renewal of the intestinal epithelium ^22,31,35,45,49,71,74–76^. However, the role of these mesenchymal cells in the maintenance and renewal of villus SM has not been explored. Gut musculature trajectories from our scRNA-seq dataset suggest that villus SM cells differentiate from local PDGFRα+ fibroblasts. We characterized mesenchymal differentiation trajectories in embryonic and juvenile mice by inducing lineage tracing in multiple pulse-chase intervals. We found that villus mesenchymal cells undergo a myofibroblast-like cell transition as they differentiate towards contractile villus SM fibers. Myofibroblast-like cells accumulate in villi during villification and begin to assemble into villus SM starting at birth. In both *Pdgfra+* and *Tnc+* cell lineage tracing, we observed *tdTomato*+ cells in a greater proportion of SM fibers at birth than during neonatal or adult lineage tracing intervals. These rates may indicate that villus SM assembly initiates at birth and then transitions into a state of regenerative turnover later in life. In the adult mouse intestine, villus mesenchymal cells are unevenly distributed along the crypt-villus axis with PDGFRα^high^ cells concentrated at the villus base ^22,35^. Additionally, TNC expression is specific to villus subepithelial fibroblasts but is absent in the crypt ^14,49^. We propose that in the highly regenerative villi, most of the subepithelial fibroblasts are primed to become TNC+ myofibroblast-like cells, ready for villus SM fiber renewal. Therefore, directing therapeutic genes toward these myofibroblast intermediates holds promise for enhancing intestinal SM repair.

Villus SM cells assemble as axial fibers, closely associated with the lacteal during perinatal development, and their interactions are critical for dietary lipid absorption ^2,5^. Villus SM is an important source of VEGF-C, the critical lymphangiogenic growth factor necessary for lacteal maintenance and lipid absorption ^12^. Genetic mutations in the embryonic mesoderm can disrupt the structure of both the villus SM and their adjacent lacteals ^24^. However, the reciprocal mechanisms guiding villus SM assembly alongside lacteals have not been described. Our work uncovers a specific upregulation of NOTCH3 signaling and a vascular SM-like gene expression signature during villus SM differentiation. DLL4 expression in lacteal tip cells is necessary for adult lacteal regeneration and lipid transport ^14^ - hence, NOTCH3 expression in the lacteal-associated villus SM suggests contact-dependent signaling within the muscular-lacteal complex. In juvenile mice, the villus muscular-lacteal complex failed to assemble correctly when *Notch3* was deleted from mesenchymal cells or when *Dll4* was removed from lymphatics. The lack of *Dll4* resulted in reduced lengths of both lacteals and villus SM, while the deletion of *Notch3* in the mesenchyme affected the length of SM only. However, both deletions led to a decreased percentage of lacteals with tip cell filopodia. These findings may suggest local Notch redundancy (and cooperation) in the intestinal villus. In our single-cell data, *Notch1* and, to some extent, *Notch2*, are both present in the villus SM cluster, consistent with a compensatory function. Finally, loss of *Notch3* resulted in impaired clearance of fats through the muscular-lacteal complex, as observed by fluorescent tracing of lipids and blood lipid measurements. These findings indicate a novel role for NOTCH3-DLL4 signaling in the assembly and lipid absorptive function of the muscular-lacteal complex. NOTCH3 plays a role in the maturation of vascular SM cells in visceral organ arteries and autonomously controls the differentiation of DLL4+ arterial endothelial cells ^56,57,61,77,77–79^. In this study, the removal of Notch3 from mesenchymal cells not only disrupted the assembly of villus SM but also impacted the coverage of mesenteric blood vascular SM cells. These observations suggest that the assembly program of villus SM shares similar characteristics with vascular SM. As a result, it is conceivable that mammalian lacteals have adopted the use of DLL4+ expression in their tip cells to facilitate the coupling and assembly of the muscular-lacteal complex. In summary, our findings reveal the molecular and cellular nature of villus SM assembly and renewal and serve as a framework for understanding the developmental specification, organization, and function of mesenchymal cell types within the intestine.

## Acknowledgments

We would like to thank P. Schweitzer and the Cornell Genomics Center for help with single-cell sequencing assays and the Cornell Bioinformatics facility for assistance with bioinformatics. Notch3 flox mice were a gift from J. Kitajewski and N. Adler; Tnc CreERT2 mice were a gift from Dr. C.M. Hao. We thank Drs. D. Gludish, P. Sethupathy, and the members of the Kurpios and De Vlaminck labs for the valuable discussion and feedback.

## Funding

National Institute of Diabetes and Digestive and Kidney Diseases R01 DK092776 (NAK); National Institute of Diabetes and Digestive and Kidney Diseases R01 DK107634 (NAK); National Institutes of Health 1DP2AI138242 (IDV); Cornell University Center of Vertebrate Genomics Scholarship (BDS, MM); Cornell University Center of Vertebrate Genomics Seed Grant (NAK, IDV); Cornell University College of Veterinary Medicine Graduate Research Fellowship (BDS); NIH 1S10RR025502 for Cornell Institute of Biotechnology.

## Author contributions

BDS, MM, IDV, and NAK designed the study. BDS and MM performed the scRNA-seq experiments and analyzed the data. BDS and MAT performed the lineage tracing experiments. BDS and LH performed the gene knockout, lipid tracing, and blood lipid measurements. BDS performed the WT and reporter immunostaining. SH and MFZW assisted with data analysis and interpretation. BDS, MM, IDV, and NAK wrote the manuscript. All authors provided feedback and comments.

## Data and code availability

The authors declare that all sequencing data supporting the findings of this study have been deposited in NCBI’s Gene Expression Omnibus with GEO series accession number <u>GSE222122</u>. All code and scripts needed to reproduce the findings of this manuscript have been deposited on GitHub: (https://github.com/madhavmantri/mouse_intestine_development). All other data supporting the findings in this study are included in the main article and associated files.

## Materials and correspondence

Requests for materials may be directed to (NAK) and (IDV).

## Conflicts

The authors declare no conflicts.

## Supplementary figure legends

**Figure S1. Single-cell RNA-seq analysis of developing mouse intestine.** Related to Figure 1. **(A)** The number of unique genes detected per cell (left), number of unique transcripts per cell (center), and percentage of mitochondrial transcripts (right) in intestinal scRNA-seq datasets from five developmental stages (E12.5, E14.5, E16.5, E18.5, and P1.5). **(B)** Bar plot showing the number of viable intestinal cell transcriptomes captured in scRNA-seq experiments. **(C)** Pie chart showing the fraction of UMIs mapping to spliced and unspliced transcripts (left) and stacked bar plot showing changes in the fraction of spliced and unspliced molecules across developmental stages (right). **(D)** Bar plot showing the proportion of cells from cell types in gut epithelium and vasculature lineages across developmental stages. **(E)** Dendrogram showing hierarchical clustering of single-cell transcriptomes from intestinal cell types across developmental stages.

**Figure S2. Single-cell RNA-seq analysis of mouse intestinal mesenchymal cells across development.** Related to Figure 2. **(A)** UMAP plot of 9,445 single-cell transcriptomes from gut musculature at 5 developmental stages (E12.5, E14.5, E16.5, E18.5, and P1.5), clustered by gene expression and colored by cluster labels. **(B)** UMAP plots showing mouse development stages for the gut musculature scRNA-seq data. **(C)** UMAP plot showing the expression of colon tissue-specific gene markers in the gut musculature scRNA-seq data. **(D)** Dot plot showing the differential gene expression analysis of mesenchymal cell types in the small intestine. Cell type labels based on UMAP in Figure 2A. **(E)** Heatmap showing the differential gene expression analysis results for three distinct smooth muscle clusters in the small intestinal scRNA-seq data. **(F)** UMAP showing *Nrp2* expression in the scRNA-seq trajectories of the small intestinal musculature and immunofluorescent staining of jejunal villi sections. NRP2 expression (on the cell membrane) is specific to the outer gut wall and mucosal lymphatic endothelium - colocalized with *Prox1*-GFP reporter (intracellular) and LYVE1 (on the cell membrane). All scale bars = 50μm.

**Figure S3. Transcription factor and gene regulatory network analysis of mouse intestinal mesenchymal cells.** Related to Figure 2. **(A)** Dot plot showing the differential transcription factor expression analysis for mesenchymal cell types in the small intestine. Cell type labels based on UMAP in Figure 2A. **(B)** Scatter plots showing Regulon Specificity Score (RSS) for transcriptional regulatory networks (regulons) for mesenchymal cell types in the small intestine. Regulon specificity scores were calculated using pySCENIC package and top five regulons for each cell type are labeled in red.

**Figure S4. Subepithelial fibroblasts and myofibroblasts in the developing gut.** Related to Figure 3. **(A)** The left image of both E18.5 and P9 panels shows whole-mount immunostaining of jejunal villi epithelium with E-CAD (on the cell membrane) and mesenchyme with PDGFRLJ (on the cell membrane), whereas the rest of the images show a single 1μm section along the z stack. PDGFRLJ is expressed in the aggregated mesenchymal cells at the villus tip and in the subepithelial lining of cells in the villus. **(B) and (C)** Immunofluorescent staining of tissue sections for PDGFRLJ and SMA (cytoplasmic). In **(B)** PDGFRLJ is broadly expressed in the proximal midgut mesenchymal cells at E12.5. e - endoderm, m - gut tube mesenchyme. In **(C)** PDGFRLJ expression was detected in SMA-subepithelial cells and was lower or absent in SMA^low^ cells SM (yellow asterisks) at E18.5. Yellow boxes represent images from the areas of the magnified insets and include DAPI+ nuclei to further appreciate cell location within the villus. All scale bars = 50μm.

**Figure S5. *Pdgfra+* lineage contribution to villus SM.** Related to Figure 3. *Pdgfra+* lineage tracing in induction-collection intervals **(A)** P8i-P9c and **(B)** P9i-P18c and P18i-27c; whole-mount immunostaining of SMA of jejunal villi. Boxes (blue and green) show the region magnified in the inset with a single 1μm section along the whole-mount z stack. Yellow arrows point to SMA+ fibers with tdTomato expression. tdT - tdTomato reporter. All scale bars = 50μm.

**Figure S6. Analyzing the heterogeneity of perinatal fibroblast-like cells from the small intestinal mesenchyme.** Related to Figure 4. **(A)** UMAP plots of 1,708 single-cell transcriptomes from perinatal fibroblast-like cells clustered by gene expression and colored by cluster labels and development stage. **(B)** Dot plot of the perinatal fibroblast-like cell clusters showing the expression of literature-derived signaling ligands and inhibitors defining adult gut fibroblast heterogeneity. **(C)** Dot plot showing the differential gene expression analysis of the perinatal fibroblast-like cell clusters. **(D)** Dot plot of the perinatal fibroblast-like cell clusters showing the expression of literature-derived genes defining adult gut fibroblast heterogeneity. **(E)** UMAP plots of the perinatal fibroblasts-like cells showing the expression of literature-derived markers associated with a myofibroblast-like cell phenotype. **(F)** UMAP plots of the perinatal fibroblasts-like cells showing the expression of Hedgehog signaling effectors in the myofibroblast-like cluster.

**Figure S7. *Tnc+* lineage contribution to villus SM**. Related to Figure 4. **(A)** Immunofluorescent staining of jejunal villi section at P9 for TNC (intra/extracellular) and SMA. **(B)** UMAPs show the expression of TNC and Pitx2, focusing on cells from E12.5 (left). Immunofluorescent staining for TNC and RNA *in situ* hybridization for *Pitx2* in a midgut section from E12.5 (right). CMA - cranial mesenteric artery, DM - dorsal mesentery, GT - gut tube. *Tnc+* lineage tracing in induction-collection intervals **(C)** P9i-P18c and P18i-27c, **(D)** P8i-P9c, and **(E)** P68i-P69; whole-mount immunostaining of SMA in jejunal villi. Boxes show the region magnified in the inset with a single 1μm section along the whole-mount z stack. In **(C)**, yellow arrows point to SMA+ fibers (cyan) with tdTomato (magenta) expression. In **(D)**, yellow arrows point to a tdTomato+ cells transitioning into SM. Yellow asterisks mark *Tnc+* lineage myofibroblast-like cells. In **(E)**, including a 3D rendering of the boxed area (right), all the magenta cells are *Tnc+* lineage myofibroblast-like cells, and there are no tdTomato+ SM fibers. tdT - tdTomato reporter. All scale bars = 50μm.

**Figure S8. Villus SM and progenitors in the human gut cell atlas.** Related to Figure 3, 4 and 5. **(A)** UMAP plot of cell clusters from the published human gut cell atlas. **(B)** UMAP plots of fibroblast and SM markers in the human gut cell atlas. Circled clusters are part of the proposed (present study) villus SM trajectory in the mouse and are pointed to by black arrows in the cell types legend. **(C)** UMAP plots of NOTCH signaling regulators enriched along the villus SM trajectory in the mouse (present study) and human gut cell atlas. Circled clusters are part of the proposed villus SM trajectory in the mouse.

**Figure S9. NOTCH-DLL4 expression in the muscular-lacteal complex.** Related to Figure 5. **(A)** UMAPs show the expression of *Wnt4* and *Acta2*, marking the cluster annotated as ‘vascular SM progenitors’ composed mainly of E14.5 cells (left). Immunofluorescent staining for WNT4 and SMA in a midgut section from E14.5 (right). Insets show WNT4+ SMA+ mesenteric vascular SM and WNT4-SMA+ gut wall SM. **(B)** Immunofluorescent staining of jejunal villi section at P9 for HEY2 (intracellular) and SMA; magenta box shows a magnified inset. **(C)** Immunofluorescent staining of jejunal villi section at P9 for MCAM (extracellular/on cell membrane) and SMA; Cyan box shows a magnified inset. MCAM is expressed in villus SM fibers and the surrounding villus blood capillaries. **(D)** Immunofluorescent staining of jejunal villi tissue sections at E18.5 for DLL4 (on the cell membrane) and LYVE1 (on the cell membrane). The Cyan arrow shows the colocalization of DLL4 and LYVE1 in the lacteal tip cell. **(E)** Immunofluorescent staining of *Prox1*-GFP (intracellular reporter) jejunal villi tissue sections at E18.5 for DLL4 (on cell membrane). A yellow dotted line outlines the lacteal with a DLL4+ tip. All scale bars = 50μm.

**Figure S10. NOTCH3 signaling regulates villus SM assembly alongside lacteals.** Related to Figure 5. **(A)** Immunofluorescent staining of jejunal villi for DLL4 and LYVE1 upon *Dll4* deletion using *Prox1-CreERT2* in the P0i-P9c interval. Cyan dotted areas highlight the lacteal tip where DLL4 expression is lost upon *Dll4* deletion. Cyan arrows highlight rare *Dll4*+ intestinal epithelial cells where *Dll4* expression was unaffected. **(B)** Immunofluorescent staining of jejunal villi for NOTCH3 and SMA upon *Notch3* deletion using *Pdgfra-CreERT2* in the P0i-P9c interval. Yellow arrows highlight villus SM where loss of NOTCH3 expression colocalized with SMA is observed upon *Notch3* deletion, whereas green arrows point to vascular SM in the sub-mucosa where NOTCH3 expression is unaffected. **(C)** Whole-mount immunofluorescent staining of jejunal villi for LYVE1 and SMA. Induced gene knockout of *Notch3* with *Hoxb6 Cre* results in a compromised assembly of villus SM alongside lacteal (left). Binned quantification of the percentage of villi with given ratios of lacteal and villus SM lengths upon *Notch3* deletion using *Hoxb6 Cre* (right). Comparison made using two-way ANOVA followed by Šídák’s multiple comparisons test (presented as mean ± SEM). iKO - inducibly knocked out. All scale bars = 50μm.

**Figure S11. Quantitative assessment of muscular-lacteal complex upon induction of *Notch3* and *Dll4* deletion.** Related to Figure 5. **(A-C)** Binned quantification of the percentage of villi with a given villus SM length normalized to villus length upon *Notch3* deletion using **(A)** *Pdgfra-CreERT2* and **(B)** *Hoxb6 Cre* drivers, and **(C)** Dll4 deletion using *Prox1-CreERT2*. **(D-F)** Binned quantification of the percentage of villi with given lacteal length normalized to villus length upon *Notch3* deletion using **(D)** *Pdgfra-CreERT2* and **(E)** *Hoxb6-Cre* drivers, and **(F)** Dll4 deletion using *Prox1-CreERT2*. Length ratios in Figure S6 A-F were compared by two-way ANOVA followed by Šídák’s multiple comparisons test (presented as mean ± SEM). **(G-I)** Quantification of the percentage of lacteals with filopodia upon *Notch3* deletion using **(G)** *Pdgfra-CreERT2*, **(H)** *Hoxb6 Cre*, and **(I)** Dll4 deletion using *Prox1-CreERT2*. The comparison was made using an unpaired t-test (presented as mean ± SEM).

**Figure S12. NOTCH3 signaling regulates SM assembly in the villus and the mesentery.** Related to Figure 5. **(A)** Whole-mount immunofluorescent staining of jejunal villi for LYVE1 and SMA. *Notch3* deletion with *Pdgfra-CreERT2* in the P8i-P9c interval (left). Binned quantification of the percentage of villi with given ratios of lacteal and villus SM lengths upon *Notch3* deletion with *Pdgfra-CreERT2* (right), compared using two-way ANOVA followed by Šídák’s multiple comparisons test (presented as mean ± SEM). **(B)** Whole-mount immunofluorescent staining of jejunal mesentery for CD31 (on the cell membrane) and SMA. *Notch3* iKO with *Hoxb6-Cre* results in loss of vascular SM coverage of the mesenteric blood vessels, whereas no changes are observed when Notch3 was deleted with *Pdgfra-CreERT2* in the P0i-P9c interval. iKO - inducibly knocked out. All scale bars = 50μm.

**Table S1.** Differential gene expression analysis results for the top 50 gene markers for all intestinal cell types in the intestinal scRNA-seq data.

**Table S2**. Differential gene expression analysis results for the top 50 gene markers for all mesenchymal cell types in the small intestinal musculature scRNA-seq data.

**Video S1. Gene expression dynamics of *Pdgfra* and *Acta2* in the villus stroma.** Related to Figure 3. 3D-rendered whole-mount image of E18.5 jejunal villi from mice with reporters *Pdgfra:H2b-GFP* (green) and *Acta2:dsRed* (magenta).

**Video S2. *Pdgfra+* lineage contributes to villus SM at neonatal stages.** Related to Figure 3. 3D-rendered whole-mount image of jejunal villi from mice with P0i-P9c *Pdgfra+* lineage tracing (magenta) and immunofluorescence for SMA (Cyan).

**Video S3. *Pdgfra+* lineage contributions to villus SM are reduced in adults.** Related to Figure 3. 3D-rendered whole-mount image of jejunal villi from mice with P60i-P69c *Pdgfra+* lineage tracing (magenta) and immunofluorescence for SMA (Cyan).

**Video S4. *Tnc+* lineage contributes to villus SM at neonatal stages.** Related to Figure 4. 3D-rendered whole-mount image of jejunal villi from mice with P0i-P9c *Tnc+* lineage tracing (magenta) and immunofluorescence for SMA (Cyan).

**Video S5. *Tnc+* lineage contributes to villus SM in adults.** Related to Figure 4. 3D-rendered whole-mount image of jejunal villi from mice with P60i-P69c *Tnc+* lineage tracing (magenta) and immunofluorescence for SMA (Cyan).

**Video S6. *Tnc-CreERT2* recombination is limited to myofibroblast-like cells at neonatal stages.** Related to Figure 4. 3D-rendered whole-mount image of jejunal villi from mice with P8i-P9c *Tnc+* lineage tracing (magenta) and immunofluorescence for SMA (Cyan).

**Video S7. *Tnc-CreERT2* recombination is limited to myofibroblast-like cells in adults.** Related to Figure 4. 3D-rendered whole-mount image of jejunal villi from mice with P68i-P69c *Tnc+* lineage tracing (magenta) and immunofluorescence for SMA (Cyan).

**Video S8. Visualizing the continuum of myofibroblast-like states during SM differentiation.** Related to Figure 4. 3D-rendered whole-mount image of jejunal villi from mice with P0i-P9c *Tnc+* lineage tracing (magenta) and reporter *Pdgfra:H2b-GFP* (green).

**Video S9. Perturbed muscular-lacteal assembly upon *Notch3* inducible knockout.** Related to Figure 5. 3D-rendered whole-mount image of jejunal villi from *Notch3 iKO; Pdgfra-CreERT2* P0i-P9c mouse with immunofluorescent staining for LYVE1 (magenta, lacteal) and SMA (Cyan, SM).

**Video S10. Stereotypical muscular-lacteal assembly in *Notch3* controls.** Related to Figure 5. 3D-rendered whole-mount image of jejunal villi from *Notch3* control P0i-P9c mouse with immunofluoresence staining for LYVE1 (magenta, lacteal) and SMA (Cyan, SM).

## Materials and methods

### Sample preparation for single-cell RNA sequencing

Timed-pregnant C57BL/6J mice were purchased from Jackson’s laboratory. We collected midguts - gut tubes extending from after the stomach to the end of the protruding midgut and the attached dorsal mesentery from E12.5 and E14.5 embryos. We collected 60 midguts at E12.5 and 35 midguts at E14.5. From the E16.5, E18.5, and P1.5 stages, we dissected open the small intestine extending from after the stomach to the end of the ileum and flushed it with ice-cold PBS. The mesentery of the duodenum was disposed of to prevent sampling of the pancreas. We sampled small intestines from 2 animals each for stages E16.5, E18.5, and P1.5. Samples were stored in cold PBS on ice for not more than 30 minutes before processing. When enough samples of a particular stage were collected, the tissue was cut into ∼1 mm pieces and transferred into a solution of Collagenase Type 1 (200 units/ml) and Hyaluronidase (100 units/ml) under mild agitation at 37°C. 5μl of the cell suspension was plated onto a hemocytometer every 5 minutes after 20 minutes of agitation and checked for the extent of dissociation. All samples reached a near single-cell suspension by the 35 - 40-minute mark and were passed through a 40 µm filter and centrifuged into a pellet. Following that, samples were resuspended in PBS containing 0.04% bovine serum albumin. Before loading the cells on 10x Chromium, Trypan Blue-stained cells from each sample were counted on an automated cell counter to determine their viability. The samples’ cell viability ranged from 83-88% after adjusting for the cell size threshold. To obtain the appropriate number of transcriptomes from viable cells (10,000 cells each for E12.5 and E14.5 stages, and 5000 each for E16.5, E18.5, and P1.5 stages), we adjusted the number of cells loaded on 10x Chromium using these cell viabilities. Sequencing libraries were prepared according to the 10x Genomics 3’ scRNA-seq library preparation protocol (v3) and were then sequenced on the NextSeq550 75bp kit.

### Single-cell RNA-seq data processing and visualization

Sequencing reads for scRNA-seq experiments were aligned to the mm10 mouse genome (assembly: GRCm38; release-93) to generate gene expression count matrices. The demultiplexing, barcode processing, gene counting, and aggregation were performed using the Cell Ranger software v3.1.0 (10x Genomics). Gene expression matrices were then read using the Scanpy (v-1.8.1) package ^80^. Transcriptomes coming from more than one cell were then labeled and removed from individual scRNA-seq datasets using the scrublet (v-0.1) package ^81^. Cells with less than 200 unique genes and genes detected in less than 10 cells were excluded from the analysis. Cells with more than 30% mitochondrial transcripts were removed as stressed/ dying cells. After quality control filtering, we analyzed a total of 37,277 single-cell transcriptomes (E12.5: 10541 cells; E14.5: 11530; E16.5: 5227 cells; E18.5: 5271 cells; P1.5: 4708 cells) across five developmental stages. The scRNA-seq data was then normalized and highly variable genes were selected with min_disp=0.5 and max_mean=3 thresholds. We then performed mean centering and scaling while regressing out total UMI counts, percent mitochondrial UMIs, S score, and G2M score, followed by principal component analysis (PCA) to reduce the dimensions of the data to the top 30 principal components. Uniform Manifold Approximation and Projection (UMAP) and the Nearest Neighbor (NN) graph were initialized in this PCA space. The cells were then clustered using the Leiden method (resolution=0.5). Differential gene expression analysis (DGEA) was performed using the rank_gene_groups function in Scanpy with the two-sided Wilcoxon statistical method. Cell-type-specific canonical gene markers along with DGEA were used to assign cell-type labels. Unsupervised clustering also gave rise to a progenitor cluster with low transcriptional diversity due to increased representation of mitochondrial genes. This cluster was simply labeled ‘mito-enriched progenitors’ and was not considered for subsequent analyses. Trajectory inference on the cell type-labeled scRNA-seq data was performed using the Partition-based graph abstraction (PAGA) method ^82^. PAGA used to initialize UMAP reductions to preserve and visualize the global topology of data at the used clustering resolutions. PAGA map was then overlaid on UMAP using the Scanpy package for visualizing the global relationships between cell types. Normalized gene expression was visualized on DotPlots, UMAP plots, and Violin plots across cell type groups. RNA velocities for single cells were calculated and visualized with the scVelo (v-0.2.4) package ^16,83^. A few cell-type clusters representing cell states of the same cell type were grouped into broad cell-type groups using cell-type-specific genes and then used for downstream analysis.

### Subclustering and analysis of intestinal mesenchymal cells and mesenchymal fibroblast cells

Normalized gene expression for all mesenchymal cell types was extracted from the combined scRNA-seq dataset. We used Scanpy to reselect the highly variable genes within that cell type group with min_disp = 0.5 and max_mean = 3 thresholds. We then performed mean centering and scaling while regressing out total UMI counts, percent mitochondrial transcripts, S score, and G2M score, followed by principal component analysis (PCA) to reduce the dimensions of the data to the top 30 principal components (PCs). Uniform Manifold Approximation and Projection (UMAP) and the Nearest Neighbor (NN) graph were initialized in this PCA space using the first 20 PCs. The cells were then reclustered using the Leiden method (resolution = 0.5) to get mesenchymal cell-type clusters. Clusters expressing colon-specific genes such as *Hoxd9*, *Hoxa9*, *Colec10*, and *Adamdec1* were excluded from the analysis. The remaining mesenchymal cells were reprocessed and clustered (resolution = 0.7) to update the mesenchymal cell type clusters. Cell-type-specific canonical gene markers along with DGEA were used to assign cell-type labels. Trajectory inference on the cell type-labeled mesenchymal scRNA-seq data was performed using the Partition-based graph abstraction (PAGA) method. PAGA graph was used to initialize UMAP reductions to preserve and visualize the global topology of data at the used clustering resolutions. PAGA map was then overlaid on UMAP using the Scanpy package for visualizing the global relationships between mesenchymal cell types. Normalized gene expression for differentially expressed genes and genes of interest was visualized on DotPlots and UMAP plots across cell-type subgroups. Normalized gene expression for the fibroblast clusters was extracted from the small intestinal mesenchymal scRNA-seq dataset. Fibroblast cells were then preprocessed and clustered (resolution = 0.4) to derive fibroblast subtype clusters. DGEA was used to assign fibroblast subtype labels.

### Animal Models

Pdgfra:H2b-GFP (JAX stock #007669) ^41^ and Acta2:dsRed (JAX stock #031159) ^42^, Rosa26 CAG-tdTomato (JAX stock #007905) ^44^, Pdgfra CreERT2 (JAX stock #032770) ^43^, Tnc CreERT2 (a gift from C. M. Hao) ^53^, Hoxb6 Cre (JAX stock #017981) ^66^, Prox1 CreERT2 (JAX stock #022075) ^67^, Notch3 flox (a gift from J. Kitajewski and N. Adler) ^64^, Dll4 flox ^65^, Prox1-GFP (RRID:MMRRC_031006-UCD) ^63^, mice were previously described. All were maintained in Cornell University’s pathogen-free barrier facility and were kept on a 12-hour light/dark cycle, with free access to standard chow pellets (Envigo) and water. All the mice used for mating were 2-6 months of age. Dams and studs were housed separately before timed mating. To track pregnancy progression, timed mated breeding pairs were checked every morning and separated upon finding a vaginal plug, staged as E 0.25. Post-natal stages are defined upon birth as P0. Sex was not taken into experimental consideration for embryonic stages and postnatal stages until P9. Only female mice were collected at stages beyond P9. Tail and toe snips collected from embryonic and postnatal mice respectively were used for genotyping by PCR amplification. For direct comparison, littermates of different genotypes were sorted into the same procedure. All animal experiments adhered to the guidelines of the Institutional Animal Care and Use Committee of Cornell University and were conducted within the scope of an approved animal protocol. Cornell University operates its animal care and uses program under the Animal Welfare Assurance on file with the Office of Laboratory Animal Welfare.

### Induction of Cre recombinase activity

Cre recombinase was activated to induce lineage tracing and gene knockout by intraperitoneal injection of tamoxifen (Sigma Aldrich, 45T5648) or 4-hydroxytamoxifen (4OHT) (Sigma Aldrich, H6278), the active metabolite of tamoxifen in peanut oil (Sigma, P2144), except for P0 pups where intragastric injections were performed ^84^. *Pdgfra CreERT2* and *Prox1 CreERT2* expressing strains were induced with a single dose of 30 mg/kg of 4OHT. This strategy was chosen as it resulted in visibly saturated reporter expression in crypt and villus mesenchymal cells and lacteals respectively and did not produce developmental defects in the WT and Cre+ mice induced in the prescribed intervals. 4OHT was suitable for this induction since *Pdgfra* and *Prox1* are genes constitutively expressed by villus fibroblast-like cells and lacteals respectively. *Tnc CreERT2* expressing strains were induced with a single dose of 100 mg/kg of tamoxifen in all the long induction periods whereas 30 mg/kg of 4OHT was used for the 1 day intervals. Tamoxifen allowed targeting of cells that express *Tnc* over a broader window to be labeled and 4OHT produced rapid recombination to label cells over a 1 day interval and neither produced developmental defects in the WT and Cre+ mice induced in the prescribed intervals.

### Mouse tissue collection, staining procedures, and image acquisition

Whole-mount and section immunostaining of embryonic, perinatal, and adult intestinal tissue was adapted from previously described protocols ^85,86^. Briefly, the first jejunal loop was dissected from mice of specified stage and genotype in ice-cold 4% paraformaldehyde (PFA)/PBS, immediately followed by PBS washes and fixed overnight at 4°C in 4% PFA/PBS or into ice-cold 100% MeOH - suitable for detecting SMA^low^ cells and removing the fluorescence from endogenous reporters. Tissues fixed in PFA were washed into PBS and stored at 4°C up to a week before whole-mount immunostaining or processed for cryosectioning by dehydrating in a gradient of sucrose/PBS solutions, embedded in OCT Compound (Tissue-Tek), and stored in −80°C until sectioning. Tissues that were fixed in 100% MeOH were stored at −20°C and rehydrated into PBS with 0.1% Tween-20, followed by a 4% PFA/PBS fixation before whole-mount immunostaining or being processed for cryosectioning as described above. All frozen sections were 15 μm thick. For immunostaining frozen sections, antigen retrieval was performed with a citrate-based antigen unmasking solution. Sections were then permeabilized with 0.1% Triton X-100 in PBS for 30 min, blocked in 3% BSA in PBS for 2 hours at room temperature, and incubated with primary antibodies diluted in blocking solution for 12-16 hours at 4°C. Sections were then washed and incubated with appropriate secondary antibodies (Invitrogen, 1:500) and DAPI (Invitrogen P36930, 1:1000, nuclear label) at room temperature for 1 hour. Samples were mounted in ProLong Gold Antifade Mountant (Thermo Fisher, P36930), and stored in the dark until imaging. For whole-mount immunostaining, tissues were blocked in 3% BSA, serum, and 0.3% Triton X-100 in PBS at 4°C for 12-16 hours, then incubated with primary antibodies 24-48 hours at 4°C. After washing for 6 hours in PBS with 0.3% Triton X-100 at room temperature, changing the wash every 30 mins, tissues were incubated in secondary antibody (1:500) for 12-16 hours at 4°C. Tissues were once again washed for 6 hours in PBS with 0.3% Triton X-100 at room temperature, changing the wash every 30 mins, and then post-fixed in 4% PFA for 12-16 hours at 4°C. Tissues were washed into PBS, sliced into 100-200 μm thick layers, and mounted with ProLong Gold Antifade Mountant. The following primary antibodies were used at a 1:100 dilution unless specified otherwise: SMA-FITC (Sigma, F3777), SMA-Cy3 (Sigma, C6198), PDGFRLJ (BD Biosciences, 558774), TNC (R&D, MAB2138), E-CAD (Invitrogen, 14324980), FOXP2 (1:250, Abcam, ab16046), NOTCH3 (Abcam, ab23426), DLL4 (R&D, AF1389), LYVE1 (Abcam, ab14917), NRP2 (R&D, AF567), HEY2 (Proteintech, 10597), MCAM (Abcam, ab75769), RFP (Rockland, 600401379), WNT4 (R&D, AF475).

Mounted samples were imaged on a Zeiss LSM 710 or LSM 880 confocal microscope using 25x, 40x or 63x objectives. To capture the entire villi, multiple adjacent fields of view were stitched together on the Zeiss Zen platform before exporting images for further analysis. Image processing, confocal image stacking, and quantifications were performed using FIJI and Imaris 9.5. 3D renderings of images were created using Imaris 9.5. *In situ* hybridization for *Pitx2* mRNA was performed on midgut sections as previously described ^51^ and images were captured on a Zeiss inverted microscope under brightfield.

### Statistical quantification

We quantified the contribution of Pdgfra and Tnc lineage to villus SM formation from confocal z-stacks of villi taken at 1μm intervals, blinded to the genotype and stage of the images. Quantifications were made for ∼15 villi from each biological replicate, with 3 biological replicates being analyzed for each lineage tracing induction interval (P0i-P9c, P8i-P9c, P9i-P18c, P18i-P27c, P60i-69c, P68i-P69c). Individual villus SM fibers from each villus were analyzed for the presence of any tdTomato+ labeling colocalizing with SMA^high^ expression. The contribution was calculated as the ratio of SMA^high^ fibers with at least one tdTomato+ cell to the total number of SMA^high^ fibers counted. Statistical analyses and graphical plotting were performed in GraphPad Prism 8 (La Jolla, CA). Comparison between different lineage tracing induction intervals was tested with one-way ANOVA and multiple comparisons with Tukey’s correction. Data were expressed as the mean of the means from each biological replicate ± standard error of the mean (SEM).

We quantified the structural changes in the villus muscular-lacteal upon inducible knockout of *Notch3* and *Dll4* from maximum intensity projections of confocal z-stacks on FIJI, blinded to the genotype of the images. Quantifications were made for 25 villi from each biological replicate, with 3 biological replicates analyzed for each of the genotypes. Lacteal lengths were measured as to the highest LYVE+ cell, not considering any filopodia that might be projecting from it. Villus SM lengths were measured as the highest point reached by an SMA^high^ fiber in a particular villus, not considering the status of other fibers in the same villus. Lacteals, villus SM, and villus lengths were consistently measured for each villus and used for internal normalizations. Statistical analyses and graphical plotting were performed in GraphPad Prism 8 (La Jolla, CA). Normalized values were sorted into bins and the data were plotted as the mean of the percentage fraction of values that fit into each bin on all the biological replicates. Cell protrusion longer than or equal to 6 μm was defined as lacteal filopodia for analysis. Comparison between genotypes was performed by two-way ANOVA with Šídák’s correction or unpaired, two-tailed Student’s t-test. Data were expressed as the mean of the means from each biological replicate ± standard error of the mean (SEM).

### *In vivo* lipid tracing

A solution of BODIPY™FL C16 (Thermo Fisher D3821) was dissolved in warmed solubilizing agent Intralipid (20% emulsion, Sigma I141) and to a final concentration reached 0.4 μg/μl. Each pup was then fed 50 μL of the reconstituted Bodipy™/Intralipid solution through a Miracle Nipple®. The pups were left with the dam for 2 hours to ensure they stayed hydrated and maintained a proper body temperature, and this timing was optimized based on when the tracer front reached the jejunal-ileal junction. After sacrificing the pups, images were taken under a Zeiss dissection microscope immediately.

### Blood lipid panel

Each pup was then fed 50 μL of warmed Olive oil (Sigma-Aldrich, O1514) solution through a Miracle Nipple®. The pups were left with the dam for 2 hours to ensure they stayed hydrated and maintained a proper body temperature as optimized previously with the tracer. Pups were sacrificed by decapitation and their blood was collected and plasma was separated by centrifugation. Samples were submitted to IDEXX bioanalytics for measuring the following parameters: triglyceride, total cholesterol, high-density lipoprotein cholesterol, and low-density lipoprotein cholesterol.

## Notes

https://www.ncbi.nlm.nih.gov/geo/query/acc.cgi?acc=GSE222122

### Competing Interest Statement

The authors have declared no competing interest.

### Summary of Updates

Much of our revision efforts focused on further detailing the cell type specificity that gives rise to the villus smooth muscle (SM). We have now expanded these analyses with multiple new experiments including quantitative lineage tracing, fluorescent reporter mice, loss-of-function analyses, functional data, and computational analysis of the publicly available human intestinal cell atlas.

