## Supplemental Figures for "Origin and adult renewal of the gut lacteal musculature from villus myofibroblasts"

Fig. S1

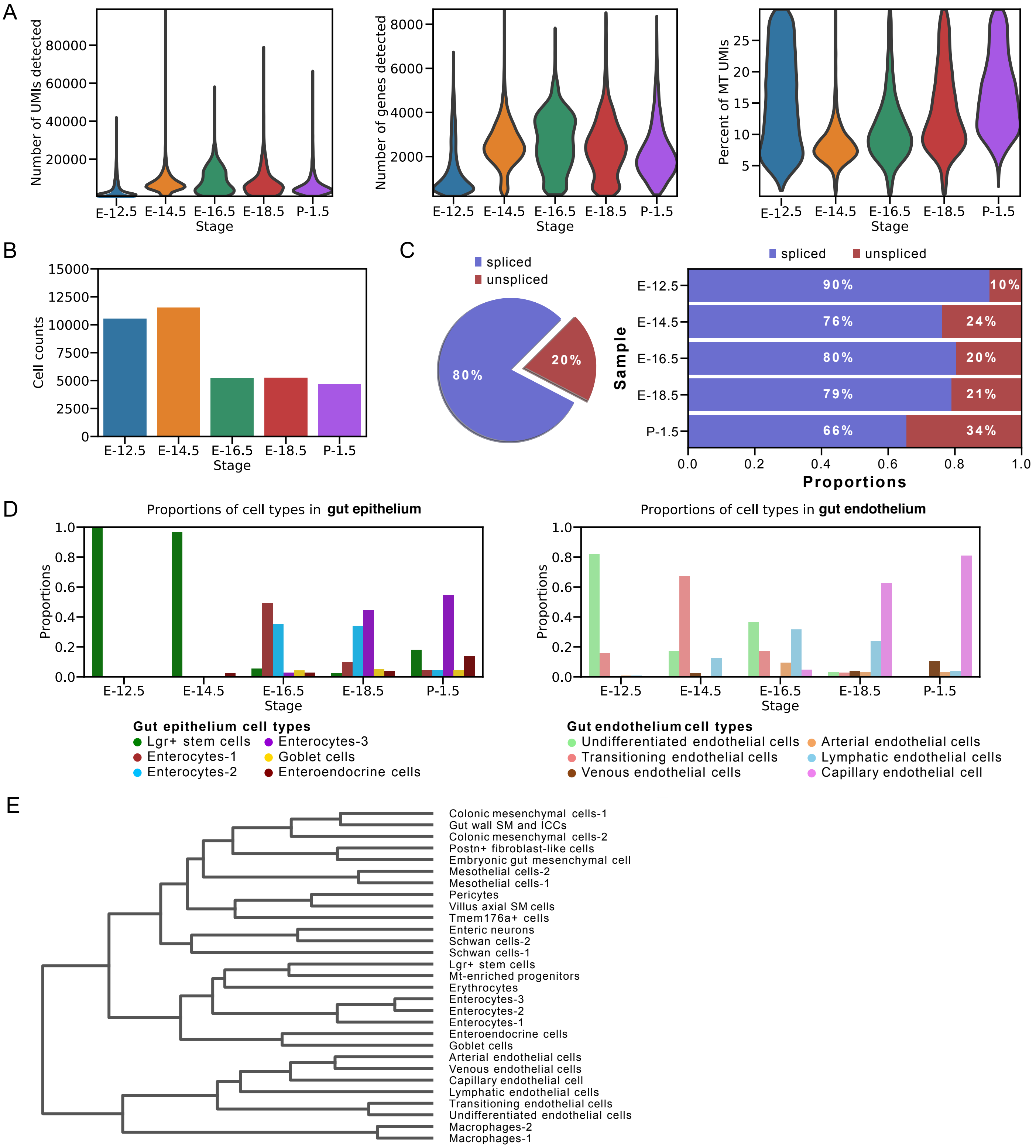
A
Number of UMIs detected
Number of genes detected
Percent of MT UMIs
Stage
E-12.5 E-14.5 E-16.5 E-18.5 P-1.5
B
Cell counts
C
spliced
unspliced
80%
20%
Sample
Proportions
E-12.5 90% 10%
E-14.5 76% 24%
E-16.5 80% 20%
E-18.5 79% 21%
P-1.5 66% 34%
D
Proportions of cell types in gut epithelium
Proportions of cell types in gut endothelium
Gut epithelium cell types
Lgr+ stem cells
Enterocytes-1
Enterocytes-2
Enterocytes-3
Goblet cells
Enteroendocrine cells
Gut endothelium cell types
Undifferentiated endothelial cells
Transitioning endothelial cells
Venous endothelial cells
Arterial endothelial cells
Lymphatic endothelial cells
Capillary endothelial cell
E
Colonic mesenchymal cells-1
Gut wall SM and ICCs
Colonic mesenchymal cells-2
Postn+ fibroblast-like cells
Embryonic gut mesenchymal cell
Mesothelial cells-2
Mesothelial cells-1
Pericytes
Villus axial SM cells
Tmem176a+ cells
Enteric neurons
Schwan cells-2
Schwan cells-1
Lgr+ stem cells
Mt-enriched progenitors
Erythrocytes
Enterocytes-3
Enterocytes-2
Enterocytes-1
Enteroendocrine cells
Goblet cells
Arterial endothelial cells
Venous endothelial cells
Capillary endothelial cell
Lymphatic endothelial cells
Transitioning endothelial cells
Undifferentiated endothelial cells
Macrophages-2
Macrophages-1

Fig. S2

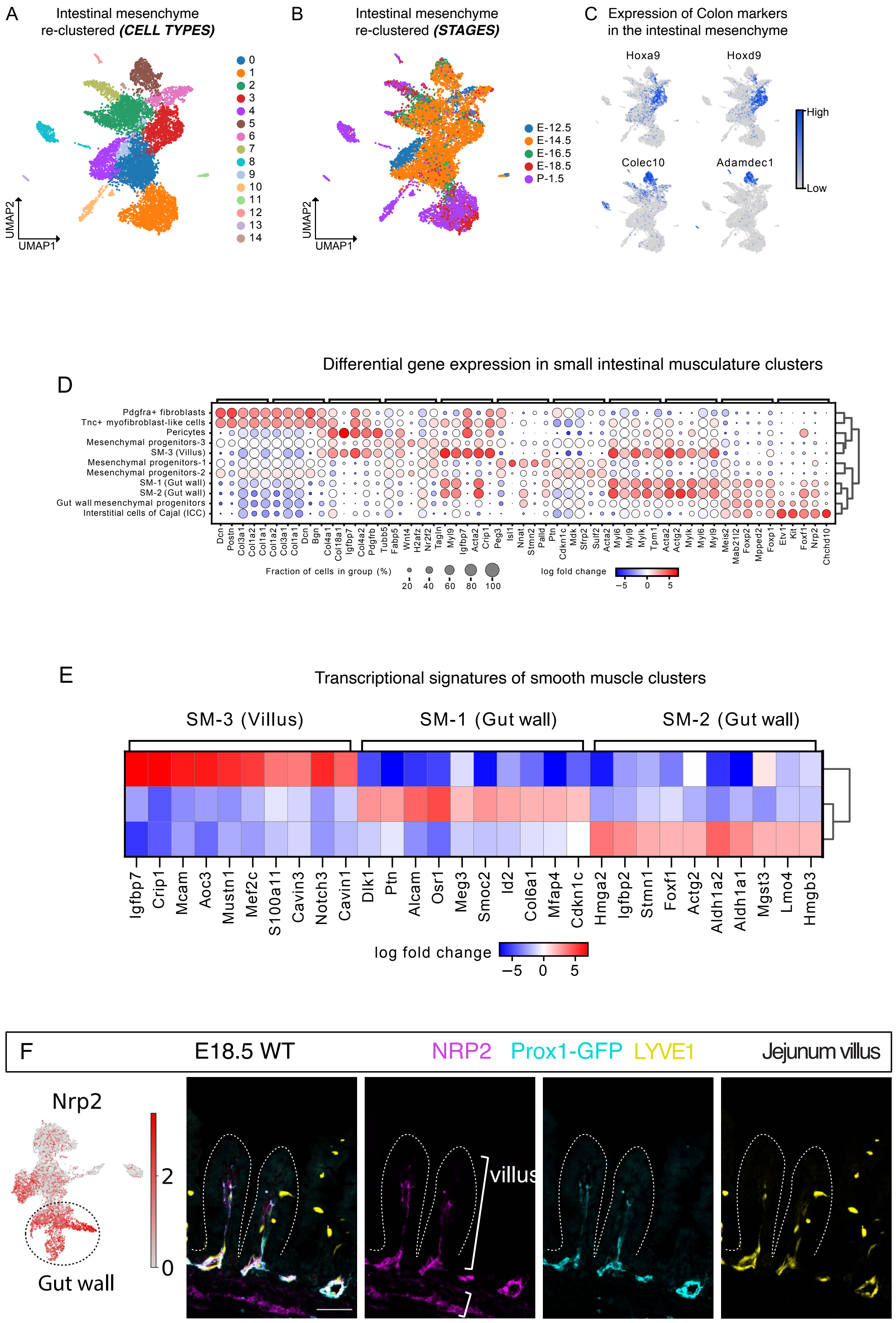
A
Intestinal mesenchyme
re-clustered (CELL TYPES)
UMAP2
UMAP1
B
Intestinal mesenchyme
re-clustered (STAGES)
E-12.5
E-14.5
E-16.5
E-18.5
P-1.5
UMAP2
UMAP1
C
Expression of Colon markers
in the intestinal mesenchyme
Hoxa9
Hoxd9
Colec10
Adamdec1
High
Low
Differential gene expression in small intestinal musculature clusters
D
Pdgfra+ fibroblasts
Tnc+ myofibroblast-like cells
Pericytes
Mesenchymal progenitors-3
SM-3 (Villus)
Mesenchymal progenitors-1
Mesenchymal progenitors-2
SM-1 (Gut wall)
SM-2 (Gut wall)
Gut wall mesenchymal progenitors
Interstitial cells of Cajal (ICC)
Dcn
Postn
Col3a1
Col1a2
Col1a1
Col1a2
Col3a1
Col1a1
Dcn
Bgn
Col4a1
Col18a1
Igfbp7
Col4a2
Pdgfrb
Tubb5
Fabp5
Wnt4
H2afz
Nr2f2
Tagln
Myl9
Igfbp7
Acta2
Crip1
Peg3
Isl1
Nnat
Stmn2
Palld
Ptn
Cdkn1c
Mdk
Sfrp2
Sulf2
Acta2
Myl6
Myl9
Mylk
Tpm1
Acta2
Actg2
Mylk
Myl6
Myl9
Meis2
Mab21l2
Foxp2
Mpped2
Foxp1
Etv1
Kit
Foxf1
Nrp2
Chchd10
Fraction of cells in group (%)
20
40
60
80
100
log fold change
−5
0
5
E
Transcriptional signatures of smooth muscle clusters
SM-3 (Villus)
SM-1 (Gut wall)
SM-2 (Gut wall)
Igfbp7
Crip1
Mcam
Aoc3
Mustn1
Mef2c
S100a11
Cavin3
Notch3
Cavin1
Dlk1
Ptn
Alcam
Osr1
Meg3
Smoc2
Id2
Col6a1
Mfap4
Cdkn1c
Hmga2
Igfbp2
Stmn1
Foxf1
Actg2
Aldh1a2
Aldh1a1
Mgst3
Lmo4
Hmgb3
log fold change
−5
0
5
F
E18.5 WT
NRP2
Prox1-GFP
LYVE1
Jejunum villus
Nrp2
2
0
Gut wall
villus

Fig. S3

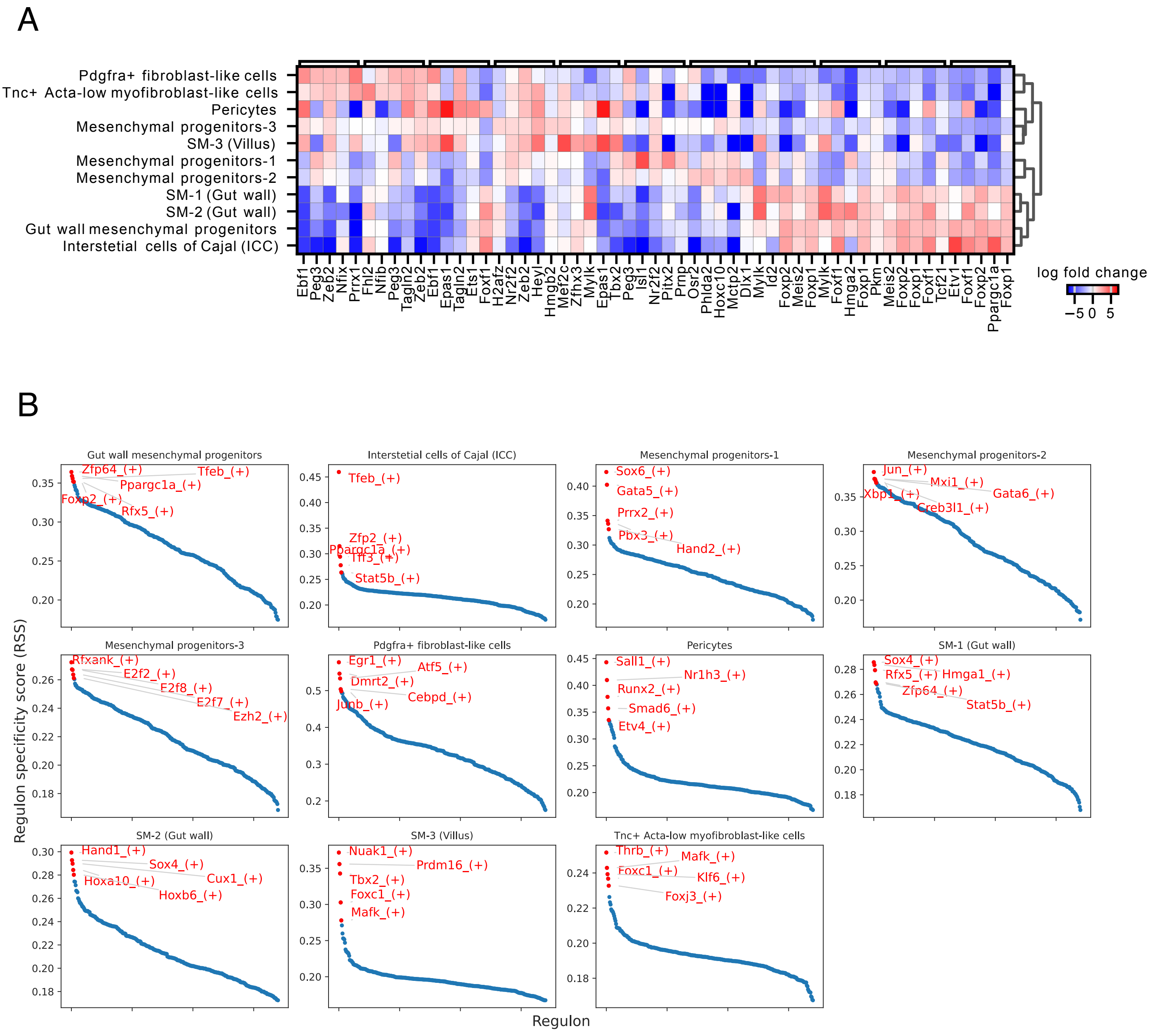
A
Pdgfra+ fibroblast-like cells
Tnc+ Acta-low myofibroblast-like cells
Pericytes
Mesenchymal progenitors-3
SM-3 (Villus)
Mesenchymal progenitors-1
Mesenchymal progenitors-2
SM-1 (Gut wall)
SM-2 (Gut wall)
Gut wall mesenchymal progenitors
Interstetial cells of Cajal (ICC)
Ebf1 Peg3 Zeb2 Nfix Prrx1 Fhl2 Nfib Peg3 Tagln2 Zeb2 Ebf1 Epas1 Tagln2 Ets1 Foxf1 H2afz Nr2f2 Zeb2 Heyl Hmgb2 Mef2c Zfhx3 Mylk Epas1 Tbx2 Peg3 Isl1 Nr2f2 Pitx2 Prnp Osr2 Phlda2 Hoxc10 Mctp2 Dlx1 Mylk Id2 Foxp2 Meis2 Foxp1 Mylk Foxf1 Hmga2 Foxp1 Pkm Meis2 Foxp2 Foxp1 Foxf1 Tcf21 Etv1 Foxf1 Foxp2 Ppargc1a Foxp1
log fold change
−5 0 5
B
Regulon specificity score (RSS)
Regulon
Gut wall mesenchymal progenitors
Zfp64_(+) Tfeb_(+) Ppargc1a_(+) Foxp2_(+) Rfx5_(+)
Interstetial cells of Cajal (ICC)
Tfeb_(+) Zfp2_(+) Ppargc1a_(+) Tff3_(+) Stat5b_(+)
Mesenchymal progenitors-1
Sox6_(+) Gata5_(+) Prrx2_(+) Pbx3_(+) Hand2_(+)
Mesenchymal progenitors-2
Jun_(+) Mxi1_(+) Gata6_(+) Xbp1_(+) Creb3l1_(+)
Mesenchymal progenitors-3
Rfxank_(+) E2f2_(+) E2f8_(+) E2f7_(+) Ezh2_(+)
Pdgfra+ fibroblast-like cells
Egr1_(+) Atf5_(+) Dmrt2_(+) Cebpd_(+) Junb_(+)
Pericytes
Sall1_(+) Nr1h3_(+) Runx2_(+) Smad6_(+) Etv4_(+)
SM-1 (Gut wall)
Sox4_(+) Rfx5_(+) Hmga1_(+) Zfp64_(+) Stat5b_(+)
SM-2 (Gut wall)
Hand1_(+) Sox4_(+) Cux1_(+) Hoxa10_(+) Hoxb6_(+)
SM-3 (Villus)
Nuak1_(+) Prdm16_(+) Tbx2_(+) Foxc1_(+) Mafk_(+)
Tnc+ Acta-low myofibroblast-like cells
Thrb_(+) Mafk_(+) Foxc1_(+) Klf6_(+) Foxj3_(+)

Fig. S4

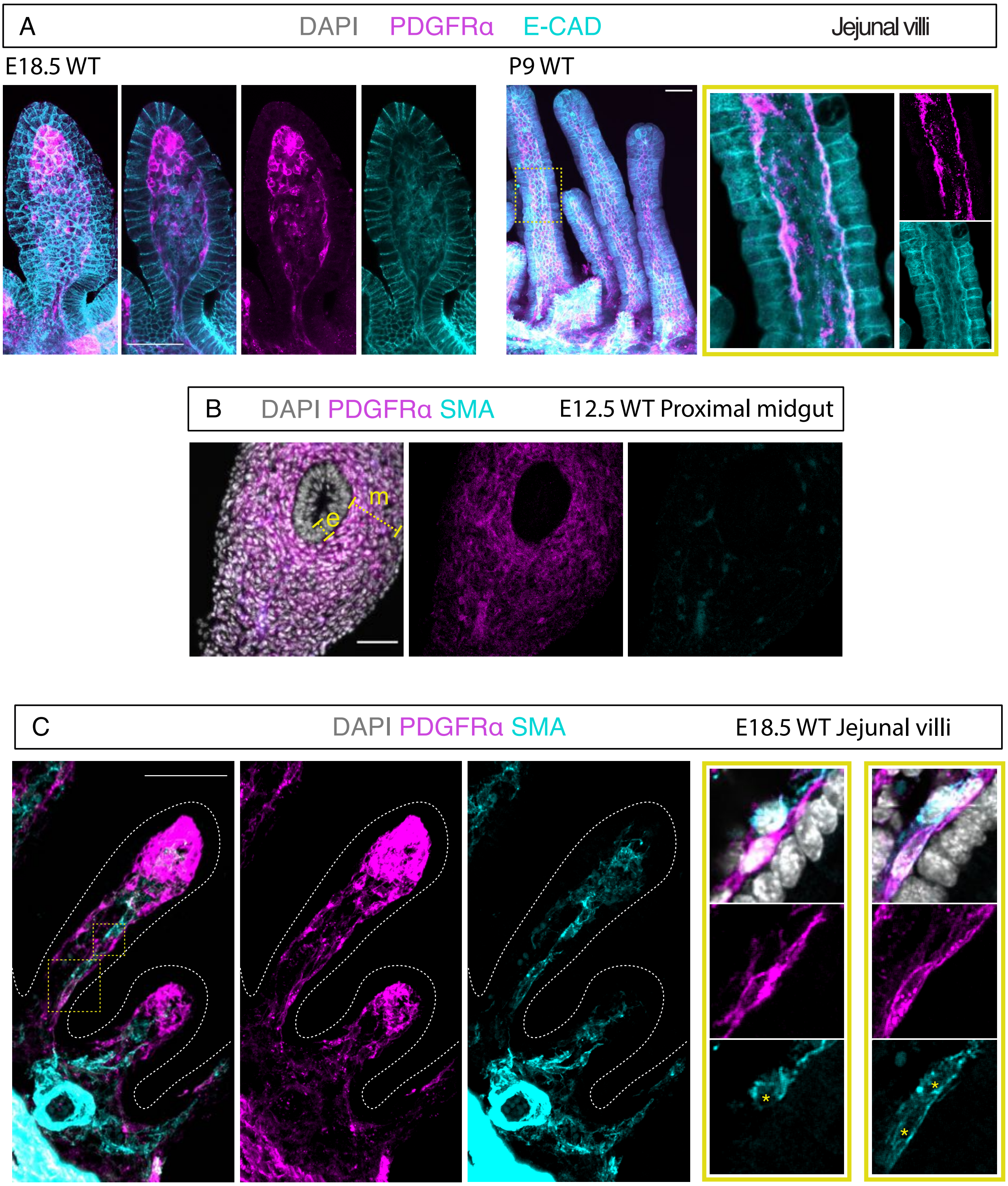
A
DAPI
PDGFRα
E-CAD
Jejunal villi
E18.5 WT
P9 WT
B
DAPI PDGFRα SMA
E12.5 WT Proximal midgut
m
e
C
DAPI PDGFRα SMA
E18.5 WT Jejunal villi
*
*
*

Fig. S5

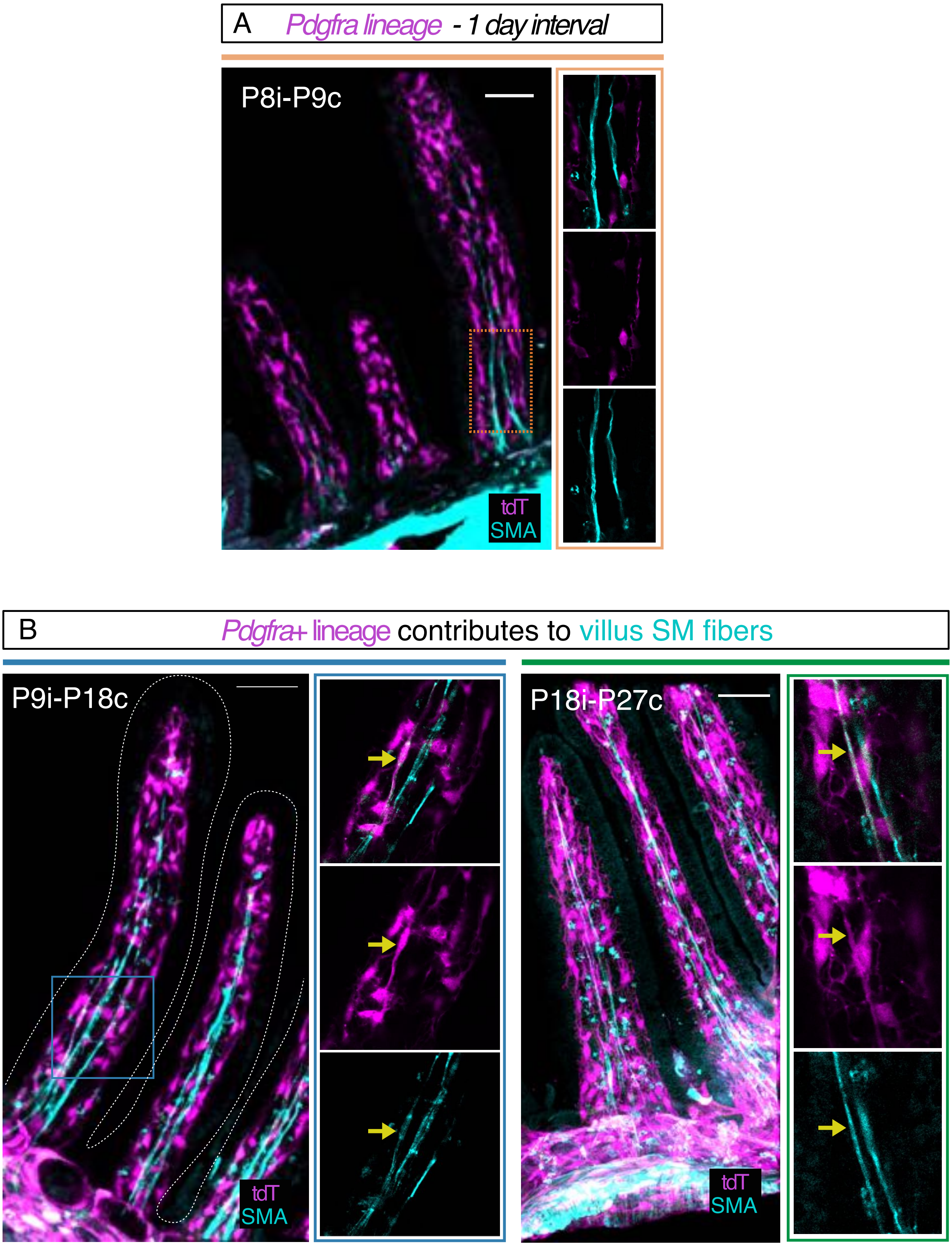
A
Pdgfra lineage - 1 day interval
P8i-P9c
tdT
SMA
B
Pdgfra+ lineage contributes to villus SM fibers
P9i-P18c
tdT
SMA
P18i-P27c
tdT
SMA

Fig. S6

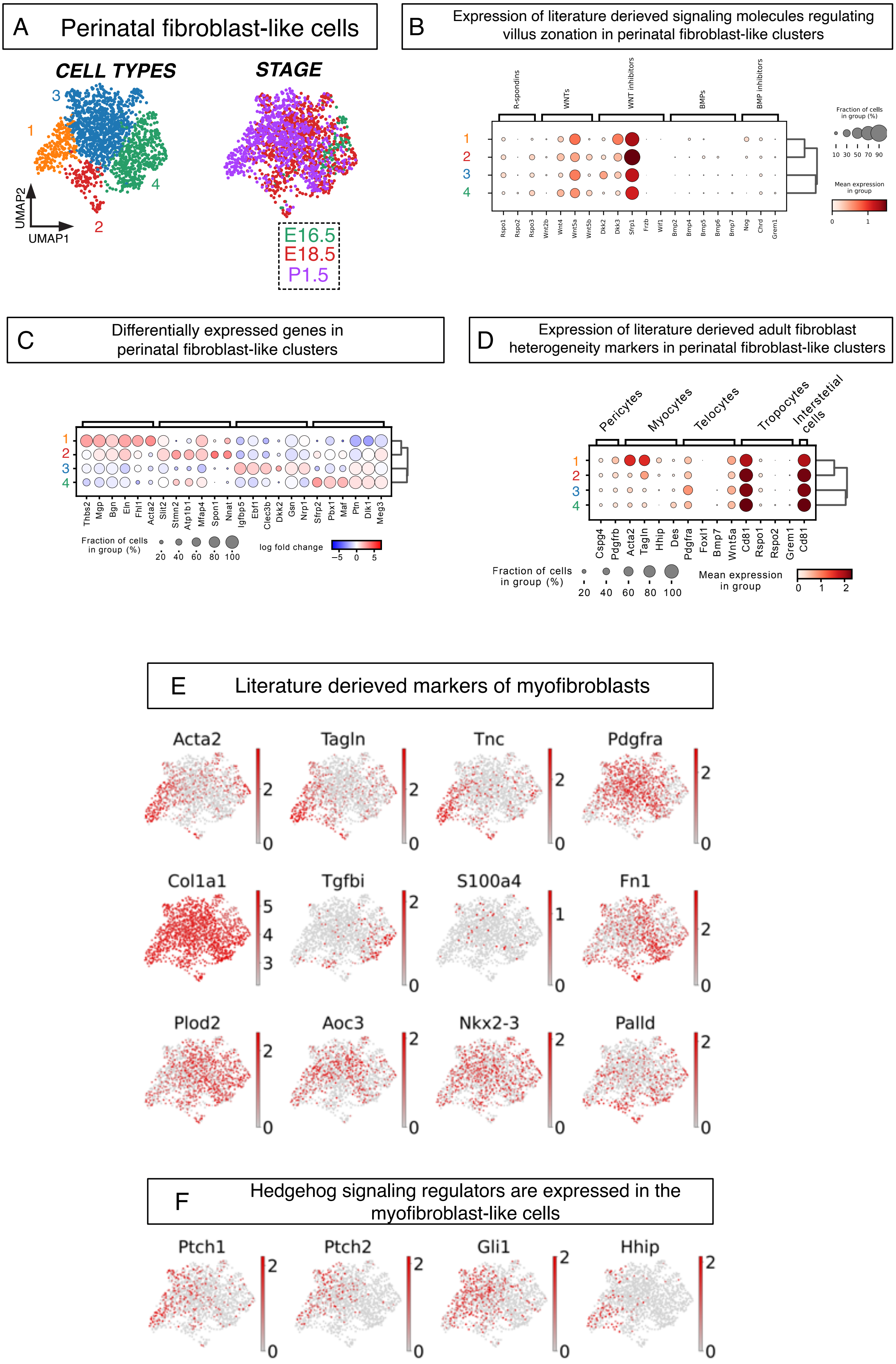

A Perinatal fibroblast-like cells
CELL TYPES
STAGE
UMAP2
UMAP1
E16.5
E18.5
P1.5
B Expression of literature derieved signaling molecules regulating villus zonation in perinatal fibroblast-like clusters
R-spondins
WNTs
WNT inhibitors
BMPs
BMP inhibitors
Rspo1 Rspo2 Rspo3 Wnt2b Wnt4 Wnt5a Wnt5b Dkk2 Dkk3 Sfrp1 Frzb Wif1 Bmp2 Bmp4 Bmp5 Bmp6 Bmp7 Nog Chrd Grem1
Fraction of cells in group (%)
10 30 50 70 90
Mean expression in group
0 1
C Differentially expressed genes in perinatal fibroblast-like clusters
Thbs2 Mgp Bgn Eln Fhl1 Acta2 Slit2 Stmn2 Atp1b1 Mfap4 Spon1 Nnat Igfbp5 Ebf1 Clec3b Dkk2 Gsn Nrp1 Sfrp2 Pbx1 Maf Ptn Dlk1 Meg3
Fraction of cells in group (%)
20 40 60 80 100
log fold change
−5 0 5
D Expression of literature derieved adult fibroblast heterogeneity markers in perinatal fibroblast-like clusters
Pericytes
Myocytes
Telocytes
Tropocytes
Interstetial cells
Cspg4 Pdgfrb Acta2 Tagln Hhip Des Pdgfra Foxl1 Bmp7 Wnt5a Cd81 Rspo1 Rspo2 Grem1 Cd81
Fraction of cells in group (%)
20 40 60 80 100
Mean expression in group
0 1 2
E Literature derieved markers of myofibroblasts
Acta2
Tagln
Tnc
Pdgfra
Col1a1
Tgfbi
S100a4
Fn1
Plod2
Aoc3
Nkx2-3
Palld
F Hedgehog signaling regulators are expressed in the myofibroblast-like cells
Ptch1
Ptch2
Gli1
Hhip

Fig. S7

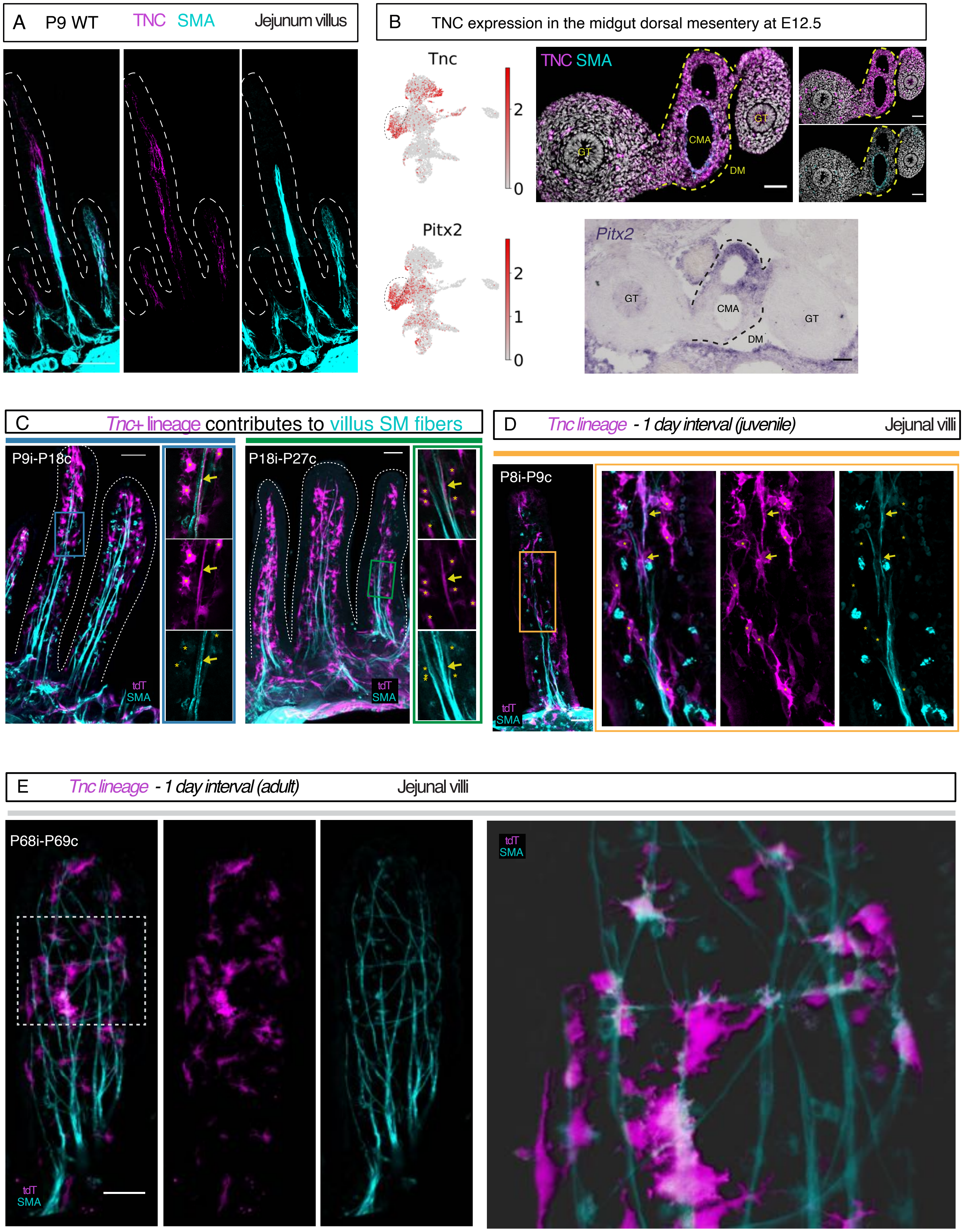

A P9 WT TNC SMA Jejunum villus
B TNC expression in the midgut dorsal mesentery at E12.5
Tnc
Pitx2
TNC SMA
GT
CMA
DM
Pitx2
C Tnc+ lineage contributes to villus SM fibers
P9i-P18c
P18i-P27c
tdT
SMA
D Tnc lineage - 1 day interval (juvenile) Jejunal villi
P8i-P9c
E Tnc lineage - 1 day interval (adult) Jejunal villi
P68i-P69c

### Fig. S8

A *Human gut cell atlas (Elmentaite et al. 2021)*

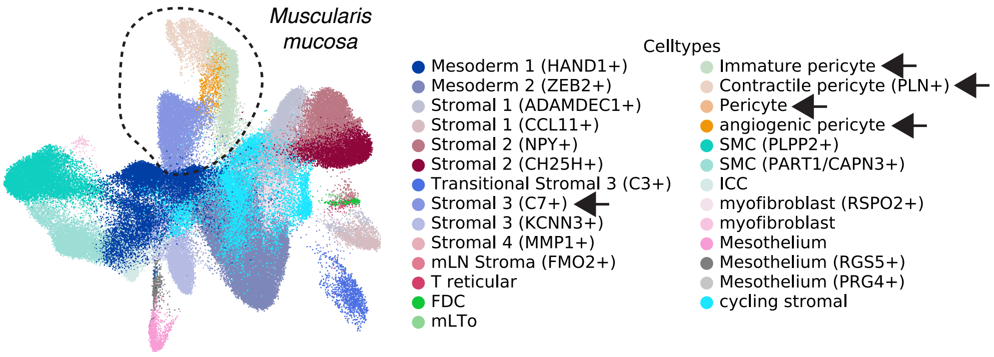

B *Fibroblast and muscle marker expression in the human gut cell atlas*

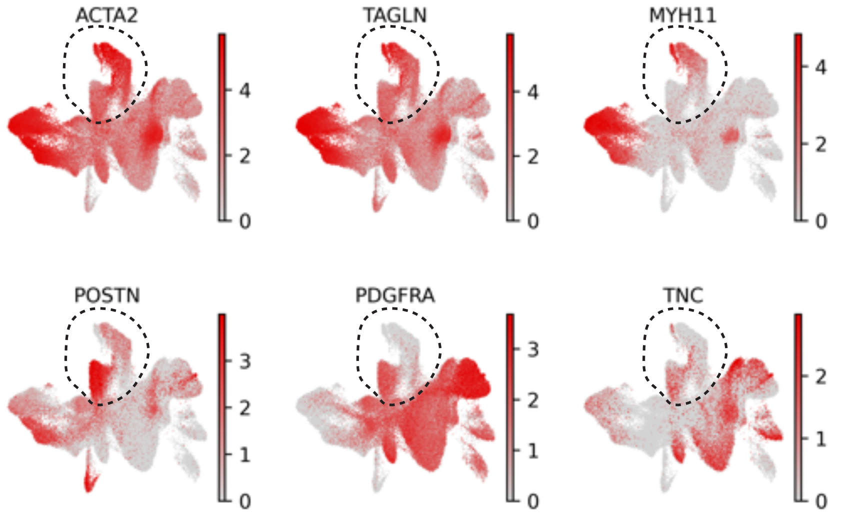

C *Conserved gene expression signature of the intestinal muscularis mucosa*

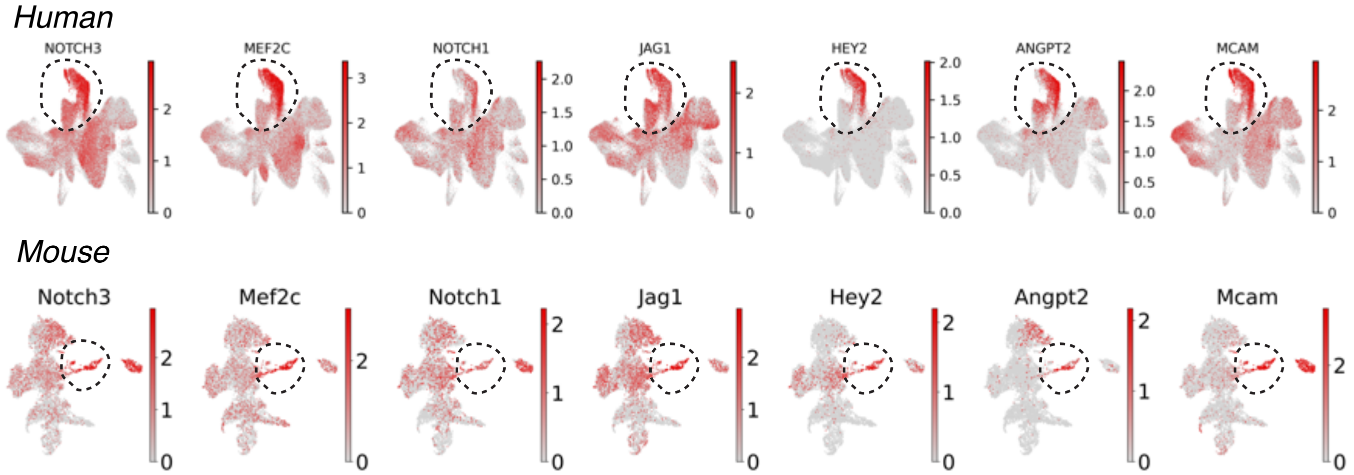

Fig. S9

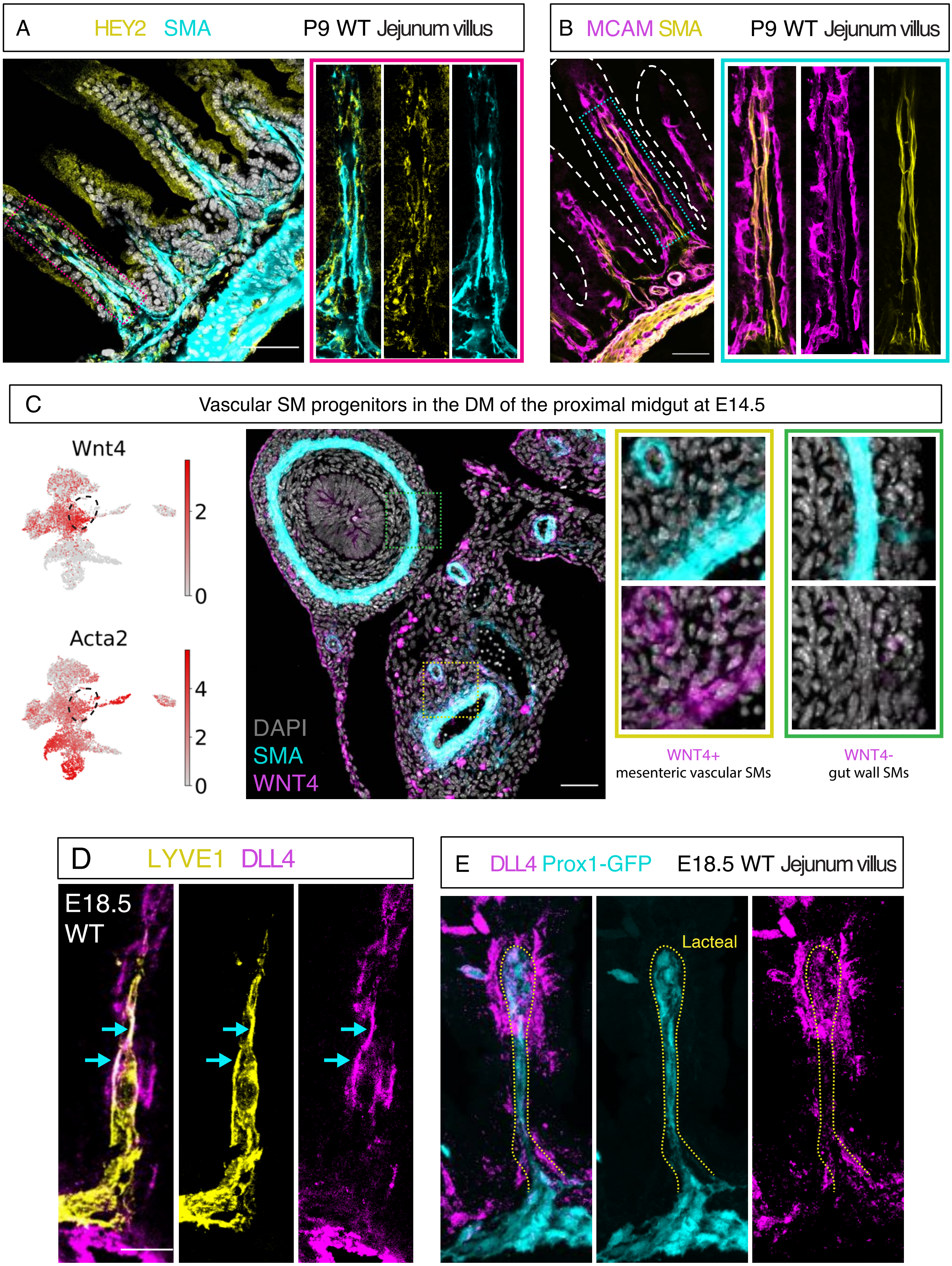

A
HEY2 SMA
P9 WT Jejunum villus
B
MCAM SMA
P9 WT Jejunum villus
C
Vascular SM progenitors in the DM of the proximal midgut at E14.5
Wnt4
2
0
Acta2
4
2
0
DAPI
SMA
WNT4
WNT4+
mesenteric vascular SMs
WNT4-
gut wall SMs
D
LYVE1 DLL4
E18.5
WT
E
DLL4 Prox1-GFP
E18.5 WT Jejunum villus
Lacteal

### Fig. S10

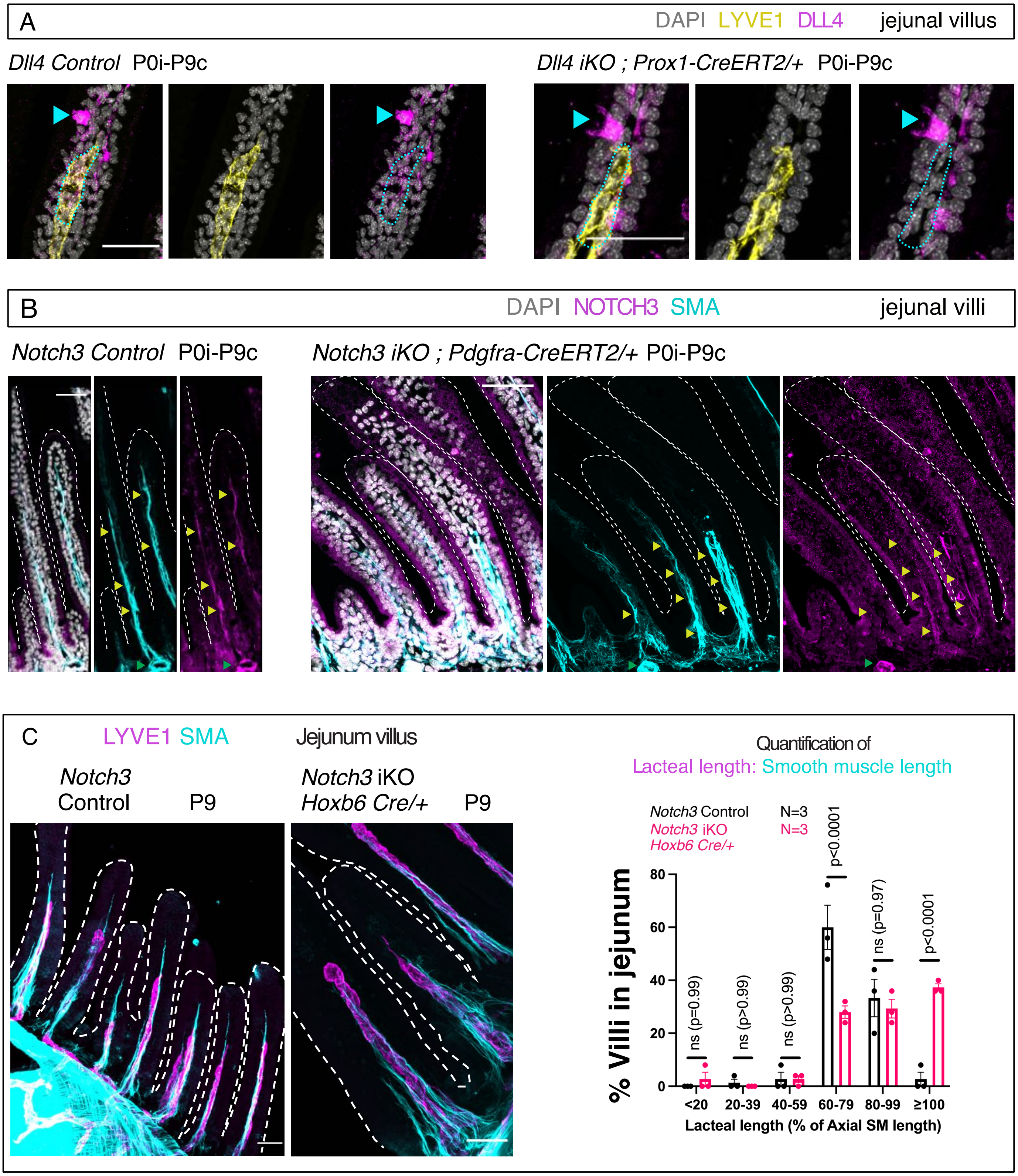
A
DAPI LYVE1 DLL4
jejunal villus
Dll4 Control P0i-P9c
Dll4 iKO ; Prox1-CreERT2/+ P0i-P9c
B
DAPI NOTCH3 SMA
jejunal villi
Notch3 Control P0i-P9c
Notch3 iKO ; Pdgfra-CreERT2/+ P0i-P9c
C
LYVE1 SMA
Jejunum villus
Notch3 Control P9
Notch3 iKO Hoxb6 Cre/+ P9
Quantification of
Lacteal length: Smooth muscle length
Notch3 Control N=3
Notch3 iKO Hoxb6 Cre/+ N=3
% Villi in jejunum
0
20
40
60
80
ns (p=0.99)
ns (p>0.99)
ns (p>0.99)
p<0.0001
ns (p=0.97)
p<0.0001
<20
20-39
40-59
60-79
80-99
≥100
Lacteal length (% of Axial SM length)

Fig. S11

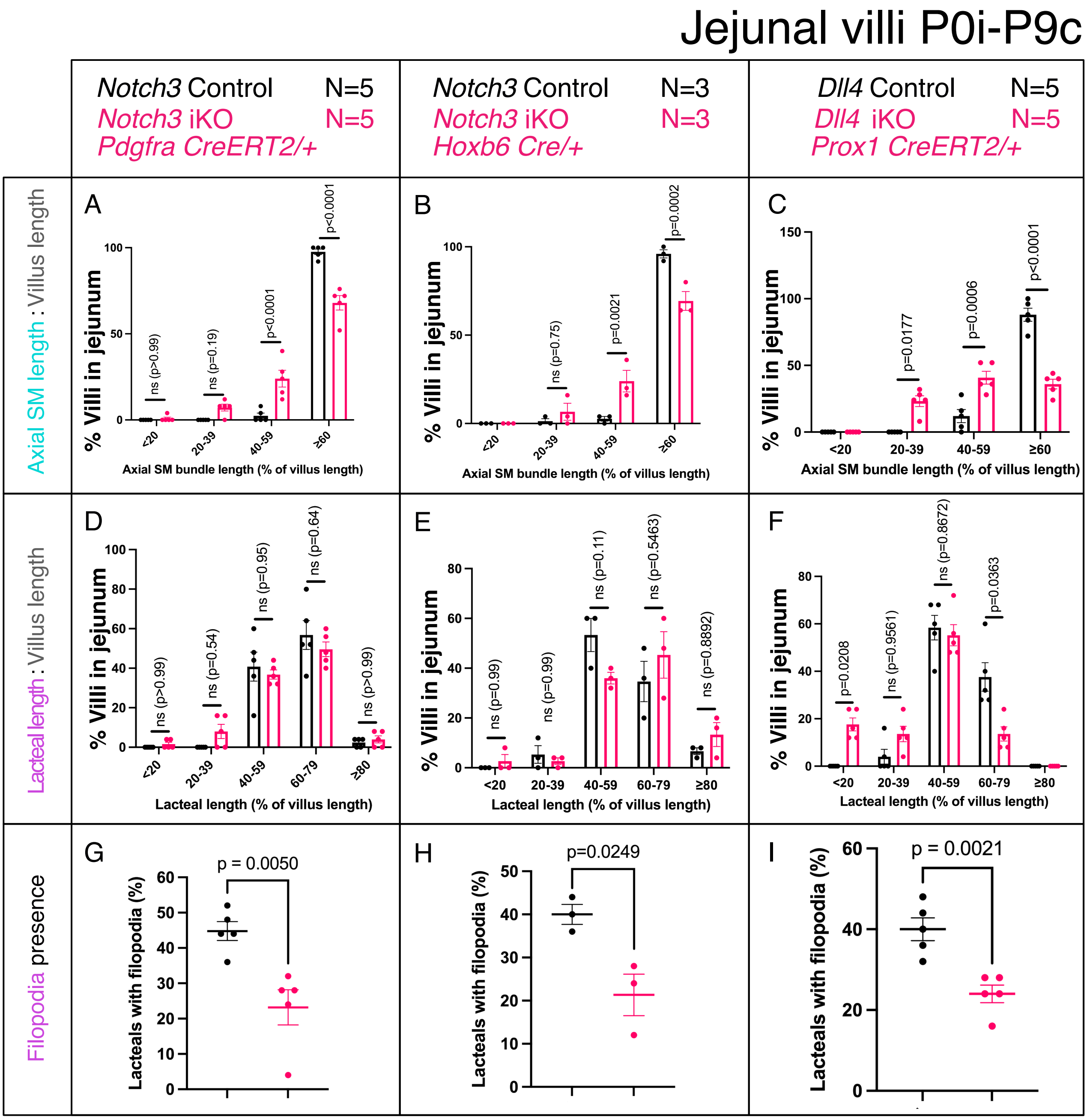

Jejunal villi P0i-P9c
Notch3 Control N=5
Notch3 iKO N=5
Pdgfra CreERT2/+
Notch3 Control N=3
Notch3 iKO N=3
Hoxb6 Cre/+
Dll4 Control N=5
Dll4 iKO N=5
Prox1 CreERT2/+
Axial SM length : Villus length
Lacteal length : Villus length
Filopodia presence
% Villi in jejunum
Axial SM bundle length (% of villus length)
Lacteal length (% of villus length)
Lacteals with filopodia (%)
A ns (p>0.99) ns (p=0.19) p<0.0001 p<0.0001
B ns (p=0.75) p=0.0021 p=0.0002
C p=0.0177 p=0.0006 p<0.0001
D ns (p>0.99) ns (p=0.54) ns (p=0.95) ns (p=0.64) ns (p>0.99)
E ns (p=0.99) ns (p=0.99) ns (p=0.11) ns (p=0.5463) ns (p=0.8892)
F p=0.0208 ns (p=0.9561) ns (p=0.8672) p=0.0363
G p = 0.0050
H p=0.0249
I p = 0.0021

### Fig. S12

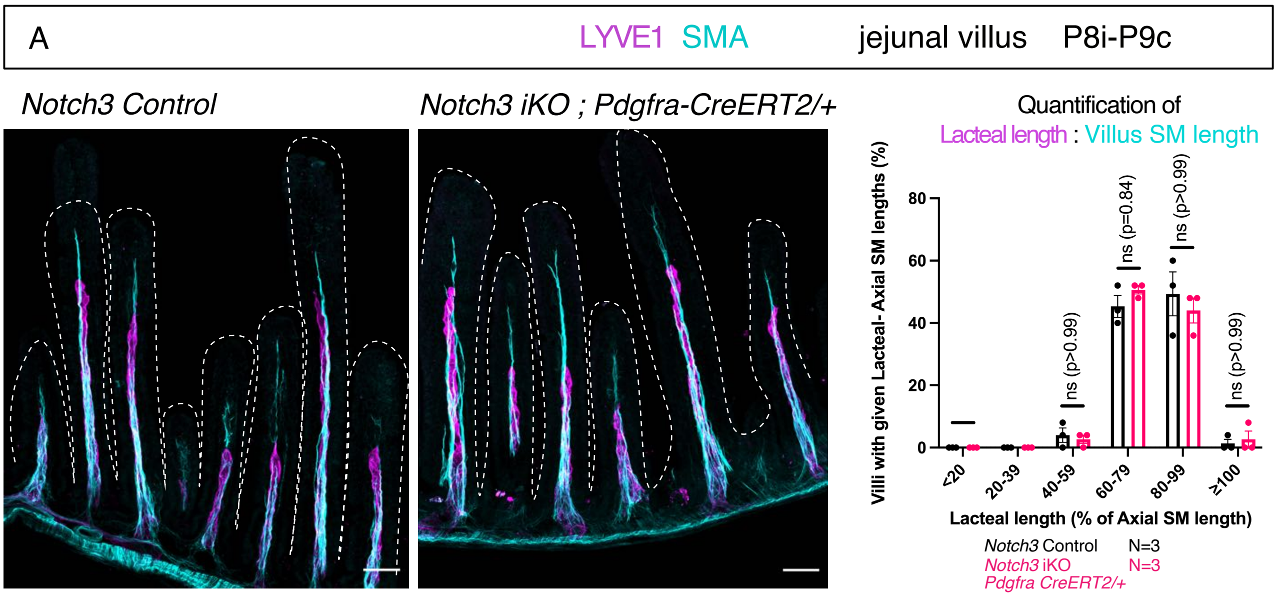
A
LYVE1 SMA
jejunal villus
P8i-P9c
Notch3 Control
Notch3 iKO ; Pdgfra-CreERT2/+
Quantification of
Lacteal length : Villus SM length
Villi with given Lacteal- Axial SM lengths (%)
80
60
40
20
0
ns (p>0.99)
ns (p=0.84)
ns (p>0.99)
ns (p>0.99)
<20
20-39
40-59
60-79
80-99
≥100
Lacteal length (% of Axial SM length)
Notch3 Control N=3
Notch3 iKO N=3
Pdgfra CreERT2/+

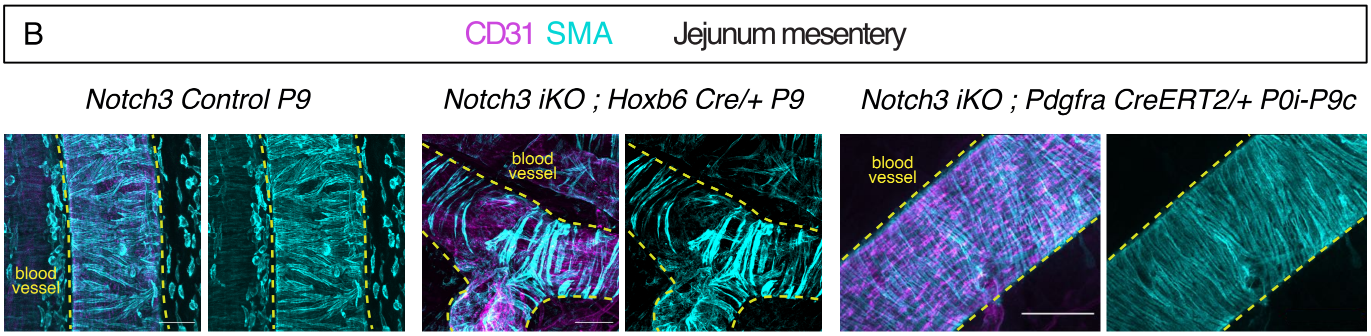
B
CD31 SMA
Jejunum mesentery
Notch3 Control P9
Notch3 iKO ; Hoxb6 Cre/+ P9
Notch3 iKO ; Pdgfra CreERT2/+ P0i-P9c
blood vessel
blood vessel
blood vessel
